# Vitamin D regulates olfactory function via dual transcriptional and mTOR-dependent translational control of synaptic proteins

**DOI:** 10.1101/2025.05.02.651619

**Authors:** Ren Pengcheng, Cao Renhe, Ye Xiaoshan, Pang Wenbin, Bi Qingshang, Huang Meihui, Zhou Qionglin, Ye Dan, Xiang Wei, Xiao Le

## Abstract

Vitamin D (VitD) deficiency, affecting over 1 billion people worldwide, is associated with neurological dysfunction, but its cell-type-specific neural mechanisms remain unclear. Using a dietary mouse model, we show that VitD bidirectionally regulates olfactory acuity: deficiency impairs odor discrimination, while supplementation enhances sensitivity. Single-nucleus and spatial transcriptomics pinpoint selective vitamin D receptor (VDR) expression in olfactory bulb (OB) tufted cells, where it drives synaptic protein expression. Genetic VDR knockdown replicates deficiency-associated olfactory deficits, establishing VDR as essential for synaptic and translational regulation. Notably, we identify mTOR-mediated protein synthesis as a critical convergence point—pharmacological mTOR inhibition (rapamycin) rescues synaptic protein deficits and behavioral impairments in VitD-deficient mice. These findings delineate a noncanonical VDR-mTOR-translational axis—complementing conventional transcriptional regulation—through which VitD serves as a nutrient-sensitive neuromodulator that integrates dietary status with synaptic functions and sensory processing. Our study expands the physiological role of VitD beyond traditional endocrine signaling and reveals mechanistic insights that may inform novel therapeutic strategies for neurological and psychiatric conditions associated with VitD deficiency.

## Introduction

Vitamin D (VitD) deficiency is a globally prevalent medical condition, affecting over 1 billion children and adults worldwide^[1]^. While traditionally recognized for its function in regulating the bone metabolism, emerging research has elucidated boarder physiological roles of VitD, particularly in the central nervous system. VitD deficiency has been increasingly associated with a spectrum of brain dysfunctions, including sensory impairment^[2]^, addiction^[3]^, depression^[4,5]^, and neurodevelopmental disorders^[6–8]^. Consequently, VitD supplementation has been investigated as both a standalone or adjunctive therapeutic strategy for these neurological and psychiatric conditions, but the efficacy of such interventions remained inconclusive^[9–13]^. Therefore, it is crucial to understand the mechanistic roles of VitD in the nervous system to fully unlock its pharmacological potential.

VitD, a fat-soluble vitamin, can cross the blood-brain barrier^[14]^, where it primarily exerts its effects by binding to vitamin D receptors (VDRs). VDRs are nuclear proteins that regulate gene expression by binding to specific DNA sequences^[15]^. In the brain, VDRs are widely distributed across regions, including olfactory cortex, cerebral cortex, hypothalamus, hippocampus and cerebellum^[16–18]^. This broad distribution highlights the potential significance of VitD in brain function but presents challenges in elucidating its region- and cell-type-specific effects. Furthermore, VitD has been implicated in the regulation of synaptic proteins, influencing key components of synaptic function such as presynaptic release machinery, neurotransmitter transporters, and receptors^[19–22]^. The expression and function of these synaptic proteins are highly context-dependent, shaped by the specific neural circuits in which they are embedded and the synaptic roles they mediate^[23]^.

Although the synaptic regulation of vitamin D, inherently linked to VDR’s function as a transcription factor, have been extensively studied^[24]^, recent attention has turned to its roles in translational control with the nervous system. For instance, a study analyzing the SFARI Gene database revealed interconnections between VitD-sensitive genes and the mechanistic target of rapamycin (mTOR) signaling pathway, an important controller of protein synthesis^[25,26]^. Additionally, phosphatase and tensin homolog (PTEN), a key regulator of mTOR signaling, has been implicated in VitD’s effects; VitD supplementation reduces social impairments in *PTEN* knockout mice^[27]^, suggesting a link between VitD and mTOR-dependent protein synthesis. However, whether VitD regulates synaptic protein synthesis through mTOR-mediated translational control, and how it is executed in a cell-type-specific manners to impact synapses and behaviors, remains unexplored.

In this study, we investigated the effects of VitD on olfactory function using a mouse model with tightly controlled VitD status from weaning to early adulthood. We demonstrated that VitD status bidirectionally regulates olfactory sensitivity. Dietary deprivation impaired odor discrimination, whereas supplementation enhanced sensory acuity. Through single-nucleus RNA sequencing (snRNA-seq) and spatial transcriptomics, we identified that the VDR is predominantly expressed in a subtype of tufted cells within the olfactory bulb (OB), where it drives cell-type-specific remodeling of dendrodendritic synapses. VitD signaling regulates synaptic and translational pathways, and VDR knockdown recapitulates olfactory impairments, as well as the synaptic and translational dysregulation observed in VitD-deficient mice. Notably, we discovered that VitD’s control over synaptic protein expression partially involves the mTOR signaling pathway. The mTOR inhibitor rapamycin rescues olfactory deficits in VitD-deficient mice by normalizing translation and excitatory synaptic protein synthesis. These findings reveal an unexpected convergence between VDR-mediated VitD signaling, transcriptional control, mTOR-dependent translational regulation, and olfactory function. The intricate interplay between molecular signaling and neuronal functions not only highlights the broader implications of VitD as a modulator of specific neural activities, extending beyond its classical endocrine roles, but also open new avenues for understanding how nutrient-sensitive signaling pathways may contribute to therapeutic applications for neurological and psychiatric disorders.

## Results

### Dietary VitD levels modulate olfactory function in mice: deficiency impairs odor discrimination while supplementation enhances sensitivity

Since VDRs are the primary mediator of vitamin D’s downstream signaling pathways, we first assessed the VDR expression at both mRNA and protein levels in the OB, cortex, hippocampus, thalamus and cerebellum. We found that the OB exhibited the highest level of VDR expression among these brain regions (Figure S1A). Furthermore, VDR expression was low at embryonic day 16 (E16) but increased postnatally, reaching consistently high levels by postnatal day 14 (P14) and persisting into two-month-old mice (Figure S1B). These findings suggest that VitD, acting through VDRs, may significantly influence olfaction from childhood through early adulthood.

Clinical observations have suggested that VitD deficiency is associated with the olfactory dysfunction, which can be alleviated by VitD supplementation in some patients^[2,28-31]^. To investigate whether varying serum VitD levels affect olfaction in mice, we employed an animal model with a controlled gradient of dietary VitD intake from childhood to early adulthood, as previously described^[19]^ (Figure 1A). In this model, mice were fed one of three diets based on AIN-93G standard chow, each containing varying dosage of vitamin D3 (VitD3), starting at weaning (P21) and continuing for approximately 11 weeks. The diets included: 0 IU/kg·day VitD3 (VitD deficiency, VDD), 500 IU/kg·day VitD3 (moderate VitD supplementation, VD-*hypo*), and 2000 IU/kg·day VitD3 (high VitD supplementation, VD-*hyper*). Serum levels of 25-(OH)D_2_ and 25-(OH)D_3_ were measured using liquid chromatography-mass spectrometry (LC-MS) (Figures S2A-S2B). The average serum 25-(OH)D levels, calculated as the sum of 25-(OH)D_2_ and 25-(OH)D_3_ levels, were significantly different between the groups (8.12±0.24 ng/ml for VDD, 22.78±1.99 ng/ml for VD-*hypo* and 53.63±2.22 ng/ml for VD-*hyper*) (Figure 1B, P<0.0001, one-way repeated ANOVA with Tukey’s multiple comparisons post hoc test). In contrast, no significant differences were observed in the average body weight, body length, or brain weight across the groups (Figure 1C, Figures S2C-S2D). Consistent with prior findings^[19]^, 25-(OH)D concentrations in the VD-*hypo* group exhibited comparable but marginally elevated levels relative to control animals maintained on standard AIN-93G diet under *ad libitum* feeding conditions (19.77±1.01 ng/ml for AIN-93G, P<0.01, unpaired t-test). Additionally, gross morphology of the OB, cerebral cortex, hippocampus and third ventricle revealed no significant changes associated with varying VitD intake (Figures S2E-S2N).

**Figure 1.**
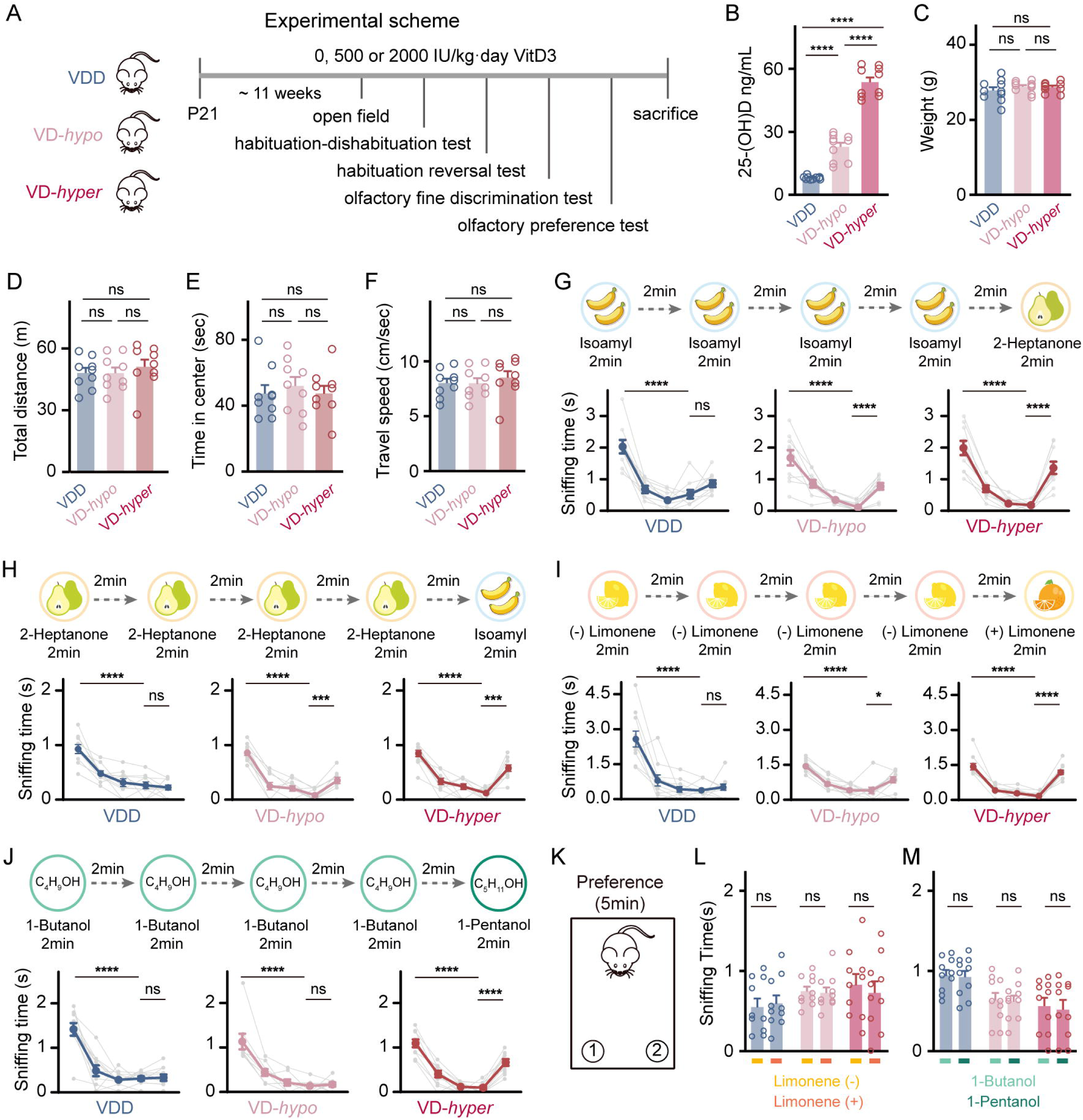
Dietary vitamin D levels bidirectionally modulate olfactory function in mice: deficiency impairs odor discrimination while supplementation enhances sensitivity. **A.** Experimental design for behavioral tests conducted on mice fed with varying doses of vitamin D3 from weaning: 0 IU/kg·day (VDD group), 500 IU/kg·day (VD-*hypo* group), and 2000 IU/kg·day (VD-*hyper* group). **B-C.** Serum 25-(OH)D levels (**B**) and body weights (**C**) of mice at 14 weeks of age (VDD, VD-*hypo*, and VD-*hyper* groups). **D-F.** Total movement distance (**D**), time to enter the middle area (**E**), and average movement speed (**F**) of mice in the open field test. **G-J.** Sniffing time in the olfactory habituation-dishabituation test (**G, I, J**) and olfactory habituation reversal test (**H**). Odor pairs were: isoamyl acetate and 2-heptanone (**G and H**), (L)-limonene and (D)-limonene (**I**), and 1-butanol and 1-pentanol (**J**). **K.** Schematic diagram of the olfactory preference test. **L-M.** Sniffing times for (L)-limonene and (D)-limonene (**L**) and 1-butanol and 1-pentanol (**M**) in the olfactory preference test. Each symbol represents a biological replicate (n=9∼10 mice per group). Data are presented as mean ± SEM. One-way ANOVA followed by Tukey’s multiple comparisons test was used for comparisons among three groups. The unpaired t-test was used for the olfactory preference test. Significance levels: *P < 0.05, **P < 0.01, ***P < 0.001, ****P < 0.0001; ns indicates no statistical significance.

Next, we conducted a series of behavioral experiments to assess basic motor function and olfactory function in the three groups of mice. In the open field test, all groups exhibited comparable performance in terms of total distance traveled, time spent in the center, and travel speed (Figures 1D-F). These findings suggest that the gradient of serum 25(OH)D levels, resulting from differential dietary VitD intake, did not affect the general motor functions in the mice.

We subsequently evaluated olfactory function using a habituation-dishabituation paradigm. Mice were initially exposed to four trials of isoamyl acetate, which emits a banana-like scent, followed by a switch to 2-heptanone, which has a pear-like scent, in the fifth trial. All groups demonstrated a significant decrease in sniffing time between the first and fourth presentations of isoamyl acetate (Figure 1G, P<0.0001, one-way repeated ANOVA with Tukey’s multiple comparisons post hoc test). However, both the VD-*hypo* and VD-*hyper* groups showed a significant increase in sniffing time upon odor renewal in the fifth trial (Figure 1G, P<0.0001, one-way repeated ANOVA with Tukey’s multiple comparisons post hoc test), whereas the VDD group did not (Figure 1G, one-way repeated ANOVA with Tukey’s multiple comparisons post hoc test). To control for potential non-specific effects of the odors, we reversed the odor presentation sequence and repeated the test. Consistent with the initial results, the VDD group again showed no significant difference in sniffing time between the fifth trial and fourth trials, while both the VD-*hypo* and VD-*hyper* groups displayed significant increases (Figure 1H, P<0.0001 for VD-*hypo* and VD-*hyper*, one-way repeated ANOVA with Tukey’s multiple comparisons post hoc test). These results indicate that VitD deficiency impairs olfactory function.

Next, we assessed the ability of the mice to discriminate between two similar odors in the habituation-dishabituation test. Mice in both the VD-*hypo* and VD-*hyper* groups showed significantly increased sniffing time at the fifth trials when the odor changed from (L)-limonene to (D)-limonene (Figure 1I, P<0.05 for VD-*hypo*, P<0.0001 for VD-*hyper*, one-way repeated ANOVA with Tukey’s multiple comparisons post hoc test). In contrast, the sniffing time of VDD mice on the fifth trials was similar to the fourth trial (Figure 1I, one-way repeated ANOVA with Tukey’s multiple comparisons post hoc test). This suggests that VitD supplementation, regardless of dosage, enabled the mice to discriminate between the similar odors of (L)-limonene to (D)-limonene.

We then increased the difficulty of discrimination by presenting two highly similar odors, 1-butanol and 1-pentanol. While all three groups exhibited habituation during the first four trials with 1-butanol (Figure 1J, P<0.0001 for VDD, VD-*hypo*, and VD-*hyper*, one-way repeated ANOVA with Tukey’s multiple comparisons post hoc test), only the VD-*hyper* group showed a significant increase in sniffing when presented with 1-pentanol on the fifth trial (Figure 1J, P<0.0001, one-way repeated ANOVA with Tukey’s multiple comparisons post hoc test). This indicates that high-dose VitD intake improves the sensitivity in odor discrimination.

As a control, we assessed the mice’s preference for the two pairs of odors ((L) - and (D)-limonene; 1-pentanol and 1-butanol) and found no significant differences between the two odors within each pair (Figures 1K-M, unpaired t-test). This confirms that the dishabituation observed on the fifth trial was due to the mice’s detection of a new odor, rather than an intrinsic preference for that odor.

In a separate cohort, we tested the olfactory behaviors of mice fed a standard AIN-93G diet *ad libitum* (Figure S3A). The performance of the standard diet group (SD group) was similar to the VD-*hypo* group. They were able to discriminate between isoamyl and 2-heptanone, as well as (L)-limonene and (D)-limonene, but not between 1-butanol and 1-pentanol (Figures S3B-S3E). Moreover, they showed similar preferences for the two pairs of odors ((L)-limonene vs. (D)-limonene, and 1-butanol vs. 1-pentanol) (Figures S3F-S3G). Notably, serum 25(OH)D levels in the standard diet (SD) group at 18 weeks (24.47 ± 0.58 ng/ml) were significantly higher than those previously observed at 15 weeks in our earlier study^[19]^ (19.77 ± 1.01 ng/ml, P < 0.01, unpaired t-test), but remained similar to levels in the VD-*hypo* group at 15 weeks (22.78 ± 1.99 ng/ml, P = 0.97). These observations suggest age-dependent fluctuations in VitD homeostasis. Furthermore, these data confirm that serum 25(OH)D concentrations and associated olfactory function in VD-*hypo* mice remain physiologically comparable to those of mice maintained on a standard diet.

In summary, using a mouse model with gradient dietary VitD intake, we found that VitD deficiency or supplementation from childhood to early adulthood did not lead to morphological changes in the OB. However, it significantly impacted olfactory function. VitD-deficient mice exhibited impairments in olfactory habituation-dishabituation and fine odor discrimination. In contrast, mice receiving VitD supplementation showed enhanced abilities in fine odor discrimination, in a dose-dependent manner.

### Layer-specific and cell-type-specific expression of VDRs in the OB

Given the significant impact of serum VitD levels on olfactory function and the high expression of VDRs in the OB, we sought to identify the specific cell types expressing VDRs in the OB. Using fluorescence *in situ* hybridization (FISH), we found VDR mRNA was most abundant in the EPL and GL, followed by the MCL, with minimal expression in the inner plexiform layer (IPL) and the granule cell layer (GCL) (Figures 2A-D). VDR-expressing (VDR^+^) neurons in the EPL and GL accounted for 79.8% of the total VDR^+^ neurons in the OB (Figure 2E).

**Figure 2.**
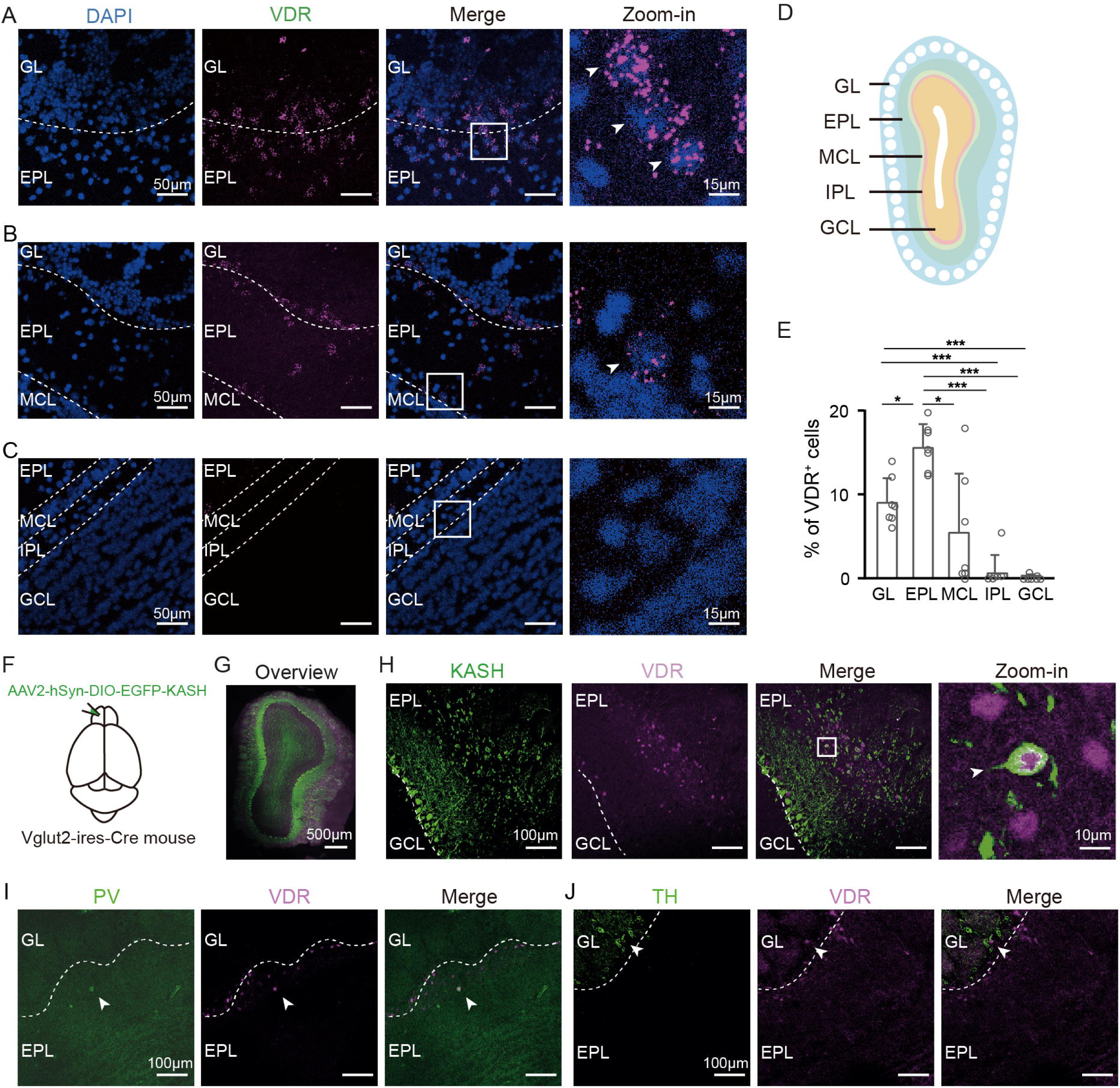
Layer-specific and cell-type-specific expression of VDR in the mouse OB. **A-C**. Representative FISH images of adult mouse OB sections showing VDR mRNA signals (purple) and nuclei counterstained with DAPI (blue). VDR expression is depicted in the GL and EPL (**A**), EPL and MCL (**B**), and IPL and GCL (**C**). Zoomed-in images highlight the framed areas, with white arrows indicating VDR mRNA signals in single cells. Scale bars: 50 µm (main images), 15 µm (zoomed-in images). **D.** Schematic drawing of the mouse OB layers, arranged from the outermost to the innermost: GL, EPL, MCL, IPL, and GCL. **E.** Quantification of VDR mRNA-positive (VDR^+^) cells across OB layers (n=3 mice). Each symbol represents a biological replicate. Data are expressed as mean ± SEM and analyzed using one-way ANOVA with Bonferroni’s multiple comparisons. Significance levels: *P < 0.05, **P < 0.01, ***P < 0.001, ****P < 0.0001. **F.** Schematic of the experimental procedure showing injection of AAV2-hSyn-DIO-EGFP-KASH-WPRES into bilateral OBs of Vglut2-ires-Cre mice, with EGFP labeling Vglut2^+^ neurons. **G.** Panoramic view of an OB section showing VDR expression (purple) and KASH (green) after viral infection and immunohistochemistry. Scale bar: 500 µm. **H.** Colocalization of KASH (green) and VDR (purple) in the EPL. The zoomed-in image (fourth column) corresponds to the framed area in the third column. Scale bars: 100 µm (main images), 10 µm (zoomed-in image). **I-J.** Immunohistochemistry showing colocalization of parvalbumin (PV, green) (**I**) or tyrosine hydroxylase (TH, green) (**J**) with VDR (purple). White arrowheads indicate neurons co-expressing marker proteins. Scale bar: 100 µm.

The EPL and GL contain excitatory tufted cells as well as inhibitory interneurons, such as periglomerular cells and short axon cells^[32,33]^. To further characterize the cell types expressing VDRs, we employed immunohistochemistry using neurotransmitter and molecular markers^[34,35]^. Since VDR is a nuclear receptor and excitatory synaptic markers are primarily localized to synapses, directly analyzing VDR protein localization relative to these markers posed challenges. To address this, we injected AAV-DIO-KASH-EGFP into the OBs of Vglut2-ires-cre mice, which express a nuclear membrane protein (KASH) and GFP in excitatory vGlut2^+^ neurons^[36,37]^ (Figures 2F-G). We found that VDRs were expressed in a subset of glutamatergic neurons in the EPL (Figure 2H). Additionally, VDR^+^ cells in the EPL and GL occasionally co-localized with parvalbumin (PV) and tyrosine hydroxylase (TH), indicating the presence of VDRs in a subset of GABAergic and dopaminergic neurons (Figures 2I-J). However, we did not observe any VDR^+^ cells co-localizing with calbindin (Calb1), calretinin (Calb2), somatostatin (SST), or S100b, suggesting that glia cells and most GABAergic interneurons do not express VDRs (Figures S4A-S4D). Together, these findings demonstrated that VDRs are present in excitatory neurons and a small population of inhibitory neurons within the EPL and GL.

### VitD signaling, mediated by VDRs, drives cell-type-specific transcriptomic remodeling about the synaptic connectivity in olfactory neurons

To unbiasedly identify the neuronal types in OB that respond to varying VitD levels and to investigate their cell-type-specific transcriptomic changes, we performed single-nucleus RNA sequencing (snRNA-seq) on OB tissues from six mice fed a gradient VitD diet (two mice per dietary group) (Figure 3A). Using snRNA-seq, we analyzed a total of 63,739 nuclei, with a median of ∼27,997 unique molecular identifiers (UMIs) and ∼1,514 genes detected per nucleus. These nuclei were annotated into 27 clusters based on established markers from the literature^[35,38,39]^. These clusters included four excitatory neuron clusters (*Slc17a6*^+^/*Slc17a7*^+^) and 18 inhibitory neuron clusters (Gad1^+^/Gad2^+^). Additionally, we identified five clusters of glia cells, comprising three astrocyte clusters (*Aqp4^+^/Gldc^+^/Emid1^+^*), one microglia cluster (*Cx3cr1^+^/Hexb^+^*), and one oligodendrocyte cluster (*Mobp^+^/Plp1^+^*) (Figures S5A-S5C). Notably, VDR expression were predominantly observed in neurons (99.35%), with only 0.65% of VDR^+^ cells identified as astrocytes (Figures S5D-S5E), consistent with our immunohistochemistry findings.

**Figure 3.**
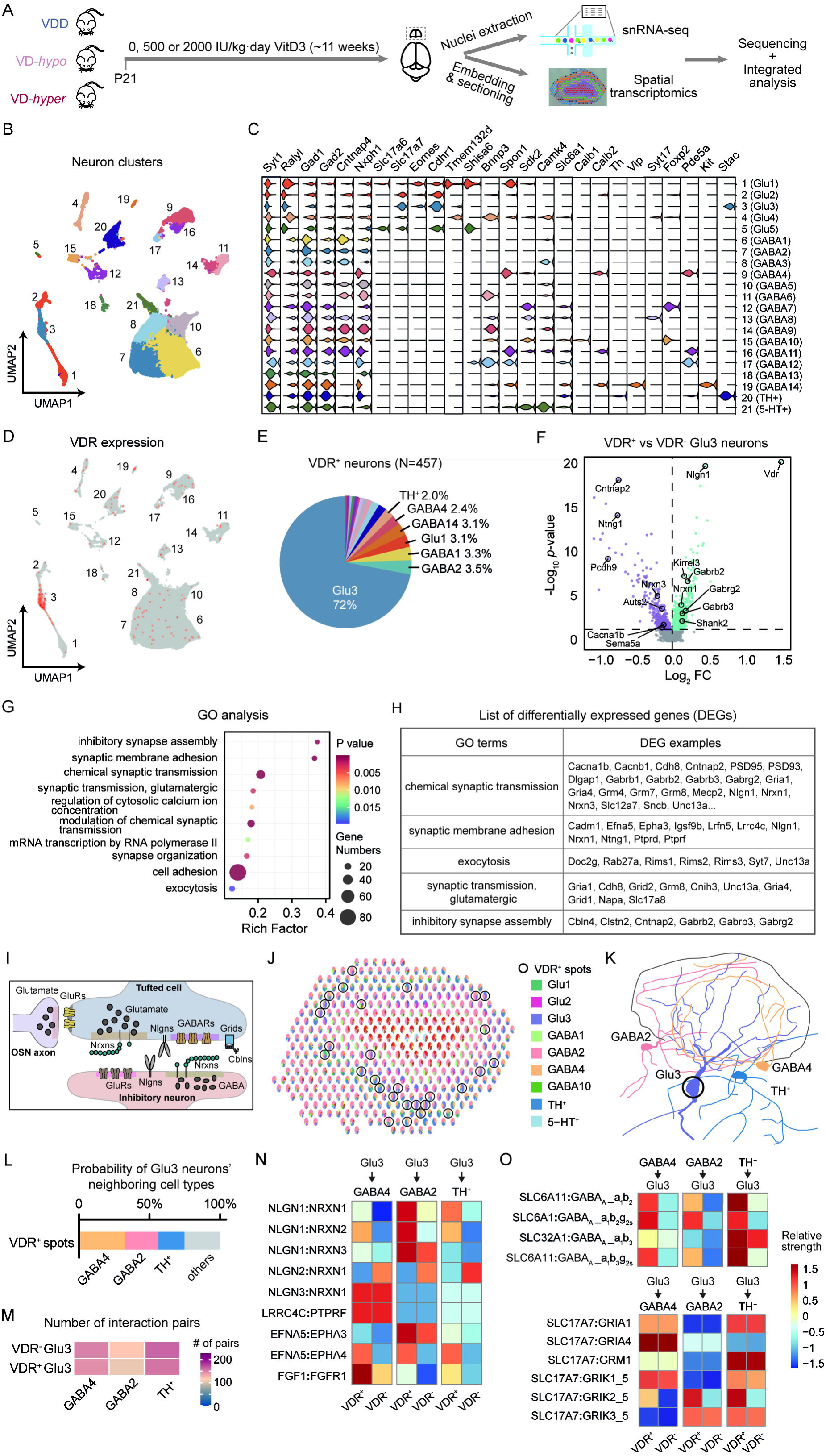
snRNA-seq and spatial transcriptomic analysis reveal transcriptomic profiles and synaptic interactions of VDR^+^ neurons in the mouse OB. **A.** Schematic flowchart of snRNA-seq (n=2 mice per group) and spatial transcriptomic analysis (n=1 mouse per group) of OBs from mice fed diets containing three different doses of vitamin D3. **B.** UMAP plot of neurons identified via snRNA-seq. **C.** Violin plots showing expression levels of selected marker genes (columns) across 21 neuronal cell clusters (rows). **D.** UMAP plot depicting the distribution of VDR-expressing neurons. **E.** Pie chart illustrating the proportion of neuron types expressing VDR. **F.** Volcano plot of DEGs between VDR^+^ and VDR^-^ Glu3 neurons. The x-axis represents log2-transformed fold change, and the y-axis shows the statistical significance (−log10 (P value)). **G.** GO annotation of DEGs between VDR^+^ and VDR^-^ Glu3 neurons. The x-axis represents the rich factor, the y-axis shows GO terms, circle size indicates the number of DEGs, and color represents P value. **H.** List of example DEGs in selected GO terms between VDR^+^ and VDR^-^ Glu3 neurons. **I.** Schematic diagram of synapses formed onto tufted cells, including dendrodendritic synapses with GABAergic interneurons and excitatory synapses with OSNs. **J.** Example image from RCTD analysis, displaying cell types in each spot from spatial transcriptomics. Black circles highlight spots expressing VDR. **K.** Schematic diagram of a Glu3 neuron and its most frequent neighboring interneurons. **L.** Probability of VDR^+^ Glu3 neurons’ neighboring cell types within a single spot from RCTD analysis. GABA4, GABA2, and TH^+^ interneurons are among the top three neighboring cell types. **M.** Heat map showing the number of interactions between Glu3 neurons and their neighboring cell types (GABA4, GABA2, or TH^+^ interneurons). **N.** Heat map of synaptic adhesion molecule interactions between VDR^+^ or VDR^-^ Glu3 neurons and their neighboring interneurons (GABA4, GABA2, or TH^+^ interneurons). **O.** Heat maps of inhibitory synaptic interactions involving GABA_A_ receptors (upper) and excitatory synaptic interactions (lower) between VDR^+^ or VDR^-^ Glu3 neurons and their neighboring interneurons (GABA4, GABA2, or TH^+^ interneurons).

Given these results, we focused on the 60,501 nuclei classified as neurons and re-analyzed them, identifying 21 clusters based on known cell-type-specific markers for excitatory projection neurons and subtype-specific markers for GABAergic interneurons from previous studies^[35,38,39]^ (Figures 3B-C). Cluster 1-5 represented glutamatergic neurons expressing *Slc17a6* (vGlut2) and/or *Slc17a7* (vGlut1). Among these, cluster 1-3 express markers characteristic of mitral/tufted (M/T) neurons, including *Eomes* and *Cdhr1^[39,40]^*, while cluster 4 and 5 represented other excitatory neuron types in the OB. Cluster 6-19 were GABAergic neurons marked by *Gad1^+^* and *Gad2^+^*. Specifically, cluster 9 (GABA4) and cluster 15 (GABA10) expressed interneuron markers (*Calb2^+^* and *Calb1^+^*, respectively) and represented two subtypes of periglomerular neurons in the GL^[41]^. Cluster 20, expressing tyrosine hydroxylase (*TH^+^*), represented short axon neurons in the GL^[41]^.

We next sought to identify the neuronal types expressing VDR. Among the 457 VDR^+^ neurons in total, 72% belonged to cluster 3 (designated as Glu3; n=155 from VDD, 103 from VD-*hypo*, 71 from VD-*hyper*), a group of excitatory projection neurons expressing vGlut1. The remaining VDR^+^ neurons included 3.5% from GABA2, 3.3% from GABA1 and 3.1% from Glu1 (Figures 3D-E). Notably, 18.45% of Glu3 neurons expressed VDRs, the highest proportion among all cell types, followed by 2.8% in GABA14 neurons and 0.63% in Glu1 (vGlut2^+^) neurons (Figure S5F, Table S1). The cell-type-specific VDR expressions observed in the snRNA-seq analysis aligns closely with our findings from FISH and immunohistochemistry.

Given that snRNA-seq analysis indicated VDRs are predominantly expressed in a specific subset of excitatory projection neurons, and FISH results localized these VDR^+^ neurons primarily to the EPL and GL, we hypothesize that VDR^+^ excitatory neurons are mainly tufted cells. These cells form dendrodendritic synapses with inhibitory interneurons, and, in some cases, receive excitatory input from olfactory sensory neurons (OSNs)^[42,43]^.

To explore cell-type-specific transcriptomic changes in the OB related to VitD, we first analyzed the most abundant VDR-expressing Glu3 neurons and identified 1,778 differentially expressed genes (DEGs) between VDR^+^ and VDR^-^ Glu3 neurons (P<0.05, log2FoldChange>1, Table S2). Notably, *VDR* itself exhibited the largest fold change between the two groups, validating our sequencing and analysis procedures (Figure 3F). Gene Ontology (GO) analysis of the DEGs revealed significant enrichment in pathways related to chemical synaptic transmission, synapse organizations and synaptic membrane adhesion (Figure 3G, Table S3). For example, genes associated with exocytosis, chemical synaptic transmission and synapse organization, such as *Rab27a*, *Unc13a*, *Cntnap2*, *Sncb*, *PSD95* (*Dlg1*), *PSD93* (*Dlg2*), *Dlgap1*, glutamate receptor delta 1 (*Grid1)*, *Cbln4*, and *Shank2,* were differentially expressed between VDR^+^ and VDR^-^ Glu3 neurons. Additionally, genes encoding glutamate receptor (*Gria1*, *Gria4*, *Grm4*, *Grm7* and *Grm8*) and GABA receptors (*Gabrb2*, *Gabrb3* and *Gabrg2*) were differentially expressed. Genes associated with synaptic membrane adhesion, such as *Nlgn1*, *Nrxn1*, *Nrxn3*, *Ntng1*, *Lrrc4c*, *Epha3*, and *Efna5*, were also differentially expressed (Figure 3H). Given that VDR^+^ Glu3 neurons are likely tufted cells based on their anatomical location and molecular markers, we found that the differentially expressed transcripts encoded synaptic proteins critical for the reciprocal dendrodendritic synapses between tufted cells and GABAergic interneurons, as well as the excitatory postsynapses formed by OSNs (Figure 3I). These findings suggest that VDR-mediated transcriptional regulation may play a pivotal role in modulating synaptic architecture within the OB, particularly in circuits involving tufted cells and their synaptic partners, which potentially shape sensory processing and integration in the OB.

We also examined VDR-associated transcriptional differences in GABAergic neurons for comparison. Due to the small number of VDR^+^ GABAergic neurons (n=91), they were analyzed collectively and compared to VDR^-^ GABAergic neurons (Figure S6A, P<0.05, log2FoldChange>1, Table S4). Interestingly, the pathways enriched in the DEGs between VDR^+^ and VDR^-^ GABAergic neurons shared some similarities but also exhibited distinct characteristics compared to Glu3 neurons. For instance, the synaptic adhesion molecule pathway (e.g., *Nlgns* and *Nrxns*), which was highly enriched in Glu3 neurons, was not significant in GABAergic neurons. Moreover, while pathways related to chemical synaptic transmission and synapse organization were enriched, the specific genes showing differential expression were distinct (Figure S6B-S6C, Table S5). For example, genes such as *Lrp4* and *Slc9a6* are differentially expressed in GABAergic neurons based on VDR expression but not in the Glu3 neurons. These findings underscore the cell-type-specific nature of VDR-mediated transcriptional regulation, and highlight the intricate and context-dependent role of VDR signaling in shaping the synaptic and functional properties of distinct neuronal populations.

We next aimed to investigate whether the neuronal interactions of Glu3 neurons, mediated by specific pairs of signaling molecules, were influenced by VDR expression. To address this, we first identified the neuronal populations spatially associated with VDR^+^ Glu3 neurons using a combination of 10X spatial transcriptomics and our snRNA-seq data (Figure 3A). The OB from one mouse per VitD dietary group (VDD, VD-*hypo*, and VD-*hyper*) was sequenced, yielding a total of 5,346 spots (55 µm per spot) in the spatial transcriptomics assay. This analysis revealed 11 distinct clusters corresponding to the six layers of the olfactory bulb (Figures S7A-S7D). Among these, 367 spots were identified as VDR^+^ (n=107 for VDD, n=61 for VD-*hypo*, n=199 for VD-*hyper*). VDR^+^ spots were predominantly classified as cluster 7 (40.3%), cluster 1 (18.8%), and cluster 2 (14.2%), which primarily correspond to the GL and EPL, corroborating our FISH data (Figure S7E).

To infer the predominant neuronal types within each spot, we employed robust cell type decomposition (RCTD) based on transcriptional profiles of major neuronal populations (defined as those with ≥ 1000 cells) derived from snRNA-seq data^[44]^. This approach ensured robust transcripts representation for accurate cell type identification (Figures S7G-S7V). For each spot, the top four cell types identified by RCTD were included in subsequent analyses. Approximately 80% of VDR^+^ spots across all dietary groups contained Glu3 neurons, consistent with our earlier snRNA-seq findings (Figure S7F).

We then focused on the VDR^+^ spots when Glu3 neurons were identified as the primary cell type by RCTD (n=177 total). Analysis of these Glu3-containing VDR^+^ spots revealed that GABA4, GABA2, and TH^+^ neurons were the most frequently observed neighboring cell types (Figures 3J). Notably, GABA4 neurons, characterized by *Calb2^+^* expression, represent a subtype of periglomerular neurons, while *TH^+^* neurons are short axon neurons. Both are inhibitory interneuron types typically localized near tufted cells in the GL and outer EPL (Figures 3L). Thus, the integration of snRNA-seq and spatial transcriptomics successfully delineated the interneuron population adjacent to Glu3 neurons.

To further explore neuronal interactions, we analyzed snRNA-seq data analyzed using Cellphone DB v5^[45]^, focusing on interactions between VDR^+^ or VDR^-^ Glu3 neurons and neighboring GABA4, GABA2, or TH^+^ interneurons (Table S6). While the overall number of interaction pairs between Glu3 neurons and these interneurons was similar regardless of VDR expressing (Figure 3M), interactions involving synaptic adhesion molecules—such as Nlgn1 with three Nrxn isoforms—were significantly stronger in VDR^+^ Glu3 neurons compared to VDR^-^ Glu3 neurons (Figure 3N). Similarly, interaction involving Efna5-Epha3/4 and FGF1-FGFR1 were predicted to be more pronounced in VDR^+^ Glu3 neurons. In contrast, interactions involving Nlgn3-Nrxn1 and LRRC4C-PTPRF showed comparable strength irrespective of VDR status (Figure 3N).

Furthermore, inhibitory synaptic interactions mediated by GABA_A_ receptors subunits (e.g., a1, b2, b3, and g2) were predicted to be stronger in VDR^+^ Glu3 neurons than in VDR^-^ neurons (Figure 3O, upper panel). Conversely, most excitatory synaptic interactions involving glutamate receptors—such as AMPA receptor subunits (Gria1 and Gria4), metabotropic glutamate receptor 1 (GRM1/mGluR1), kainate receptor subunit (e.g., Grik1/5 and Grik3/5)—exhibited similar strength regardless of VDR expression. However, the interaction involving kainate receptor subunit Grik2/5 was notably stronger in VDR^+^ Glu3 neurons compared to VDR^-^ Glu3 neurons (Figure 3O, lower panel).

In summary, the integration of Cellphone DB analysis with snRNA-seq and spatial transcriptomics data revealed that VDR^+^ Glu3 neurons—primarily tufted cells—engage in more robust and nuanced interactions with neighboring periglomerular and short axon cells within the glomerular layer. These interactions are particularly enriched for synaptic adhesion molecules, including Nlgn-Nrxn pairs, as well as specific subtypes of glutamate and GABA_A_ receptors, which collectively suggest sophisticated regulations on synaptic interactions underpinned by VDR-mediated VitD signaling.

### VitD modulates synaptic pathways in the OB via VDR-dependent mechanisms

We next examined the dose-dependent effect of VitD mediated by VDRs. To ensure a relatively homogenous cellular components in VDR^+^ spots, and to be enable comparability with VDR^+^ Glu3 neurons located in the GL and outer EPL, we selected Glu3-containing spots within the same anatomical layers corresponding to cluster 1, 2, and 7. Comparative analyses between the VD-*hypo* and VDD groups, as well as between the VD-*hyper* and VD-*hypo* groups, revealed that DEGs in VDR^+^ Glu3 neurons and Glu3-containing VDR^+^ spots were significantly enriched in pathways associated with synaptic functions, including synaptic organization and chemical synaptic transmission (Figure 4A, P<0.05, log2FoldChange>1, Table S7 and S8). Additionally, Glu3-containing VDR^+^ spots in both comparison groups, as well as VDR^+^ Glu3 neurons in the VD-*hypo* vs. VDD comparison, exhibited significant enrichment in pathways related to synaptic vesicle exocytosis and synaptic membrane adhesion. Notably, pathways involving translation and the regulation of translation were prominently enriched in VDR^+^ Glu3 neurons and Glu3-containing VDR^+^ spots under varying VitD levels. These pathways were less pronounced in comparison between VDR^+^ and VDR^-^ Glu3 neurons, suggesting that VitD exerts a specific and dose-dependent influence on translational processes in VDR-expressing Glu3 neurons, potentially contributing to the functional changes of OB.

**Figure 4.**
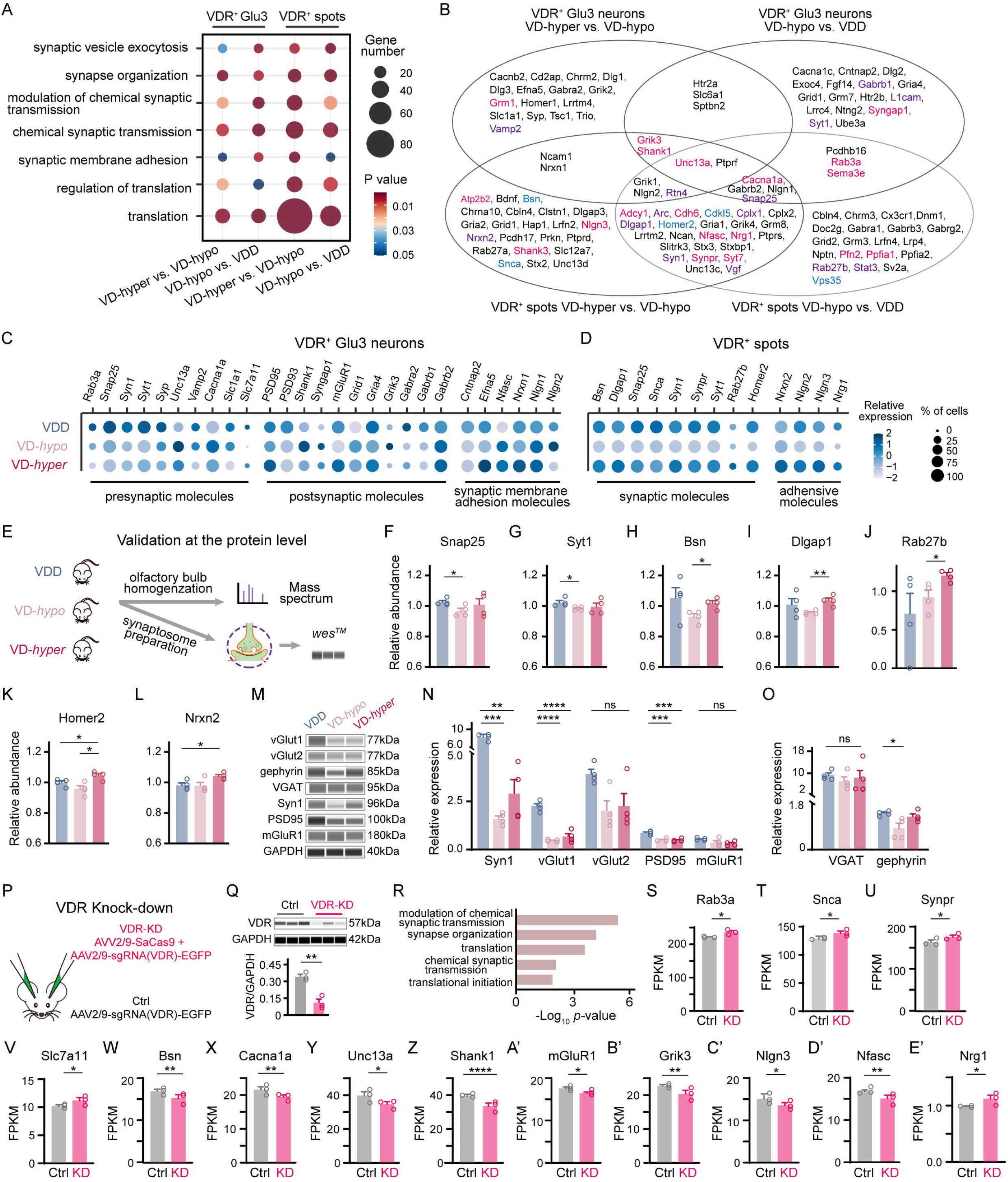
Vitamin D modulates synaptic pathways in the OB via VDR-dependent mechanisms. **A.** GO annotation of DEGs between VD-*hyper* vs. VD-*hypo* and VD-*hypo* vs. VDD in the VDR□ Glu3 group from snRNA-seq (n=2 mice per group; n=155 for VDD, n=103 for VD-*hypo*, n=71 for VD-*hyper*) and VDR□ spots from spatial transcriptomics (n=1 mouse per group; n=35 for VDD, n=22 for VD-*hypo*, n=115 for VD-*hyper*). The y-axis represents GO terms, circle size indicates the number of DEGs, and color represents P value. **B.** Venn diagram showing example overlapping DEGs from snRNA-seq (comparing VDR□ Glu3 groups: VD-*hyper* vs. VD-*hypo* and VD-*hypo* vs. VDD) and spatial transcriptomics (comparing VDR□ spots: VD-*hyper* vs. VD-*hypo* and VD-*hypo* vs. VDD). Genes marked in pink represent DEGs confirmed by RNA sequencing in control vs. VDR knockdown (VDR-KD) mice; genes in purple correspond to DEPs from proteomic analysis (VD-*hyper* vs. VD-*hypo* or VD-*hypo* vs. VDD); genes in blue are confirmed by both techniques. **C-D.** Dot plots showing expression of synaptic function-related DEGs in VDR□ Glu3 neurons from snRNA-seq (**C**) and VDR□ spots from spatial transcriptomics (**D**) across three groups (VD-*hyper*, VD-*hypo*, and VDD). Dot size represents the percentage of cells expressing the gene; color indicates averaged scaled expression levels. **E.** Experimental schematic of protein-level validations, including LC-MS-based proteomic analysis and quantitative protein analysis of whole OB tissues and synaptosome preparations using *wes* in three mouse groups (VDD, VD-*hypo*, VD-*hyper*). **F-L.** Relative expression levels of *Snap25* (**F**), *Syt1* (**G**), *Bsn* (**H**), *Dlgap1* (**I**), *Rab27b* (**J**), *Homer2* (**K**), and *Nrxn2* (**L**) in LC-MS-based proteomic analysis (n=4 mice per group; one-way ANOVA with Tukey’s multiple comparisons). **M-O.** Excitatory (**M, N**) and inhibitory (**M, O**) synaptic protein levels in synaptosome preparations from OBs of three mouse groups, measured by *wes* (n=4 mice per group; one-way ANOVA with Tukey’s multiple comparisons, except for vGlut2, analyzed using the Mann-Whitney U test due to non-normal distribution). **P.** Experimental schematic of CRISPR-Cas9-mediated VDR-KD in the mouse OB. **Q.** The *wes* image and quantification of VDR expression in VDR-KD and control (Ctrl) groups (n=3 mice per group). **R.** GO annotation of DEGs between VDR-KD and control groups (n=3 mice per group). The x-axis represents −log10(P value), and the y-axis shows GO terms. **S-E’.** Relative mRNA expression of *Rab3a* (**S**), *Snca* (**T**), *Synpr* (**U**), *Slc7a11* (**V**), *Bsn* (**W**), *Cacna1a* (**X**), *Unc13a* (**Y**), *Shank1* (**Z**), *mGluR1* (**A’**), *Grik3* (**B’**), *Nlgn3* (**C’**), *Nfasc* (**D’**), and *Nrg1* (**E’**) in control (Ctrl) and VDR-KD mice, analyzed by RNA sequencing (n=3 mice per group). Each symbol represents a biological replicate. Data are presented as mean ± SEM. Significance levels: *P < 0.05, **P < 0.01, ***P < 0.001, ****P < 0.0001; ns indicates no statistical significance. FPKM: Fragments per kilobase million.

Furthermore, we analyzed DEGs in VDR^+^ GABAergic neurons under varying VitD conditions. Similar to Glu3 neurons, translation emerged as one of the top enriched pathways in GABAergic neurons. However, pathways related to synapse organization and chemical synaptic transmission were less prominent in GABAergic neurons, indicating a degree of cell-type-specific regulation in VitD signaling (Figure S6D-S6F, Table S9 and S10). Collectively, these findings underscore shared patterns across spatial and single-nucleus transcriptomic datasets, highlighting translation and synaptic functions as key physiological domains significantly modulate by VitD levels. Although the snRNA-seq and spatial transcriptomic analyses were performed with cell-type or spot-type specificity, bulk RNA sequencing of the OB from VDD, VD-*hypo* and VD-*hyper* mice further corroborated that molecules associated with synaptic functions and translations were strongly influenced by serum VitD levels (Figure S8A). Gene set enrichment analysis (GSEA) of DEGs between the VD-hyper and VD-*hypo* groups, as well as the VD-*hyper* and VDD groups (P<0.05, log2FoldChange>1), revealed significant enrichment in pathways such as regulation of neurotransmitter receptor activity, maintenance of synapses structures, olfactory transduction, ribosome function (Figure S8B-S8D). Moreover, GO analysis of the DEGs between these two comparisons revealed significant enrichment in pathways such as chemical synaptic transmissions, cytoplasmic translation, and negative regulation of TORC1 signaling—a pathway implicated in translation regulation (Figure S8E-S8F, Table S11 and S12). These results demonstrate that varying levels VitD intake exert significant effects on synaptic functions and translational processes within the OBs, consistent with the snRNA-seq and spatial transcriptomics results.

We further looked into genes associated with synaptic functions that were differentially expressed in VDR^+^ Glu3 neurons and/or VDR^+^ Glu3-containing spots upon varying VitD levels (Figure 4B-D). Transcripts encoding presynaptic molecules, such as *Rab3a*, *Snap25*, *Synapsin1* (*syn1*), *Synaptagmin1* (*syt1*), *Synaptophysin* (*syp*), *Vesicle-Associated Membrane Protein 2 (Vamp2)*, and the glutamate transporters *Slc1a1* and *Slc7a11*, were significantly upregulated in VDR^+^ Glu3 neurons from the VDD group compared to the supplemented groups. In contrast, only two presynaptic molecules, *Unc13a* and *Cacna1a* (a voltage-gated calcium channel), were downregulated in VDR^+^ Glu3 neurons from the VDD groups (Figure 4C). Transcripts encoding postsynaptic molecules exhibited more complexed expression patterns. For instance, genes encoding postsynaptic density components, such as *PSD95* and *PSD93*, were mostly highly expressed in VDR^+^ Glu3 neurons from the VDD group, whereas *shank1* and *syngap1* showed the lowest expression in these neurons under VitD deficiency (Figure 4C). Among glutamate receptor transcripts, *Grik3* was most highly expressed, while *mGluR1* and *Gria4* were least expressed in the VD-*hypo* group, and *Grid1* was least expressed in VDR^+^ Glu3 neurons from the VDD group (Figure 4C). Additionally, transcripts for synaptic membrane adhesion molecules, such as *Cntnap2*, *Efna5*, *Neurofascin (Nfasc)*, *Nrxn1*, *Nlgn1* and *Nlgn2*, displayed altered expression in VDR^+^ Glu3 neurons under varying serum VitD levels (Figure 4C).

Interestingly, in VDR^+^ spots, many synaptic molecules, including *Bassoon* (*Bsn*), *Dlgap1*, *Snap25*, *Snca*, *Syn1*, *Synaptoporin* (*Synpr*), *Syt1* and *Homer2*, as well as synaptic membrane adhesion molecules like *Nlgn2*, *Nlgn3*, and *Neuregulin 1* (*Nrg1)*, exhibited higher expression in the VDD and VD-*hyper* groups compared to the VD-*hypo* group. Conversely, *Rab27b* and *Nrxn2* were the least expressed in VDR^+^ spots from the VDD group and most highly expressed in the VD-*hyper* group. In summary, these findings establish that differential VitD levels modulate synaptic protein expression specifically in VDR-expressing Glu3 neurons and discrete VDR-positive spots, revealing a cell-type-specific molecular mechanism through which dietary VitD intake regulates olfactory synapses.

To validate the VitD-dependent changes in synaptic molecules at the protein level, we employed two complementary approaches (Figure 4E). First, we conducted quantitative proteomic analysis using LC-MS to identify differentially expressed proteins (DEPs) in the OB of VDD, VD-*hypo* and VD-*hyper* groups (P<0.05, log2FoldChange>1, Table S13). Synaptic proteins such as Snap25 and Syt1 were significant upregulated in the VDD group compared to the VD-*hypo* group (Figures 4F-G, P<0.05, log2FoldChange>1, unpaired t-test), while Bsn and Dlgap1 were significant downregulated in the VD-*hypo* group compared to VD-*hyper* group (Figures 4H-I, P<0.05 for Bsn, P<0.01 for Dlgap1, unpaired t-test). Additionally, Rab27b, Homer2 and Nrxn2 exhibited the highest expression in the VD-*hyper* group (Figure 4J-L, P<0.05, unpaired t-test).

Second, we quantified synaptic proteins in the synaptosome isolated from the OB of mice fed with gradient VitD diets. Excitatory presynaptic proteins, including Syn1 and vGlut1, as well as the postsynaptic protein PSD95, were significantly upregulated in the VDD group compared to the supplemented groups (Figures 4M-N, one-way repeated ANOVA with Tukey’s multiple comparisons post hoc test). Similarly, vGlut2 and mGluR1 showed a trend toward increased expression in the VDD group compared to the VD-*hypo* and VD-*hyper* groups, although these changes did not reach statistical significance (Figure 4M-N). Furthermore, the inhibitory postsynaptic protein gephyrin was significantly upregulated in the VDD group, while the inhibitory presynaptic proteins VGAT remained unchanged across different VitD levels (Figure 4M, 4O).

In summary, the protein-level findings for synaptic molecules were consistent with the results from snRNA-seq and spatial transcriptomics. Together, these results demonstrate a pronounced upregulation of excitatory synaptic proteins and a moderate upregulation of inhibitory synaptic proteins in the OB under VitD deficiency. These molecular changes may provide a potential mechanism underlying behavioral alterations in olfactory function associated with varying VitD intake.

We next sought to determine whether the observed changes in synaptic protein expression in the OB under gradient serum VitD levels were mediated by the VDR. To investigate this, we knocked down VDR expression in the OB of C57BL/6 mice using adeno-associated virus (AAV) vectors encoding SaCas9 and VDR-specific single-guide RNA (sgRNA) (Figure 4P). This approach achieved a reduction of VDR protein levels by over 60% in the knock-down (KD) group compared to controls (Figure 4O). Bulk RNA sequencing followed by GO analysis of DEGs revealed significant enrichment in pathways related to synaptic functions and translation, consistent with our earlier results (Figure 4R, P<0.05, log2FoldChange>1, Table S14 and S15).

Notably, transcripts encoding presynaptic molecules—*Rab3a*, *Snca*, *Synpr*, and *Slc7a11*—were significantly upregulated in the KD group, while *Bsn*, *Cacna1a* and *Unc13a* were significantly downregulated (Figure 4S-Y, P<0.05 for *Rab3a*, *Snca*, *Synpr*, *Slc7a11* and *Unc13a*; P<0.01 for *Bsn* and *Cacna1a*). Similarly, postsynaptic molecules such as *Shank1*, *mGluR1* and *Grik3*, as well as synaptic membrane adhesion molecules like *Nlgn3* and *Nfasc*, showed significant downregulation in the KD group (Figure 4Z-D’, P<0.05 for *mGluR1* and *Nlgn3*; P<0.01 for *Grik3* and *Nfasc*; P<0.0001 for *Shank1*). In contrast, *Nrg1* was significantly upregulated (Figure 4E’, P<0.05). These results closely aligned with our sequencing and protein-level analyses, further underscoring the critical role of VDR in regulating synaptic protein expression in the OB.

### VitD regulates translation via transcriptional control, as well as VDR-mediated mTOR signaling in olfactory neurons and synapses

Given our snRNA-seq, spatial transcriptomics, bulk RNA sequencing and VDR knockdown analyses collectively indicated a strong regulation on translation by VitD levels, we next investigated the expression of translation-related genes in VDR^+^ Glu3 neurons and VDR^+^ spots under varying VitD conditions (Figure 5A-C). Most DEGs encoding translation initiation factors—such as *eIF3D*, *eIF3E*, *eIF3H*, *eIF5*—as well as the elongation factor *eEF1A1* and ribosome proteins (*Rpl10*, *Rpl18* and *Rps13*), exhibited the lowest expression in VDR^+^ Glu3 neurons from the VD-*hyper* group (Figure 5B). In contrast, *eIF2S1* and *Rpl5* showed the highest expression in the VDD group, while *Rpl7*, *Rpl37* and *Rpl37a* were most highly expressed in the VD-*hypo* group (Figure 5B).

**Figure 5.**
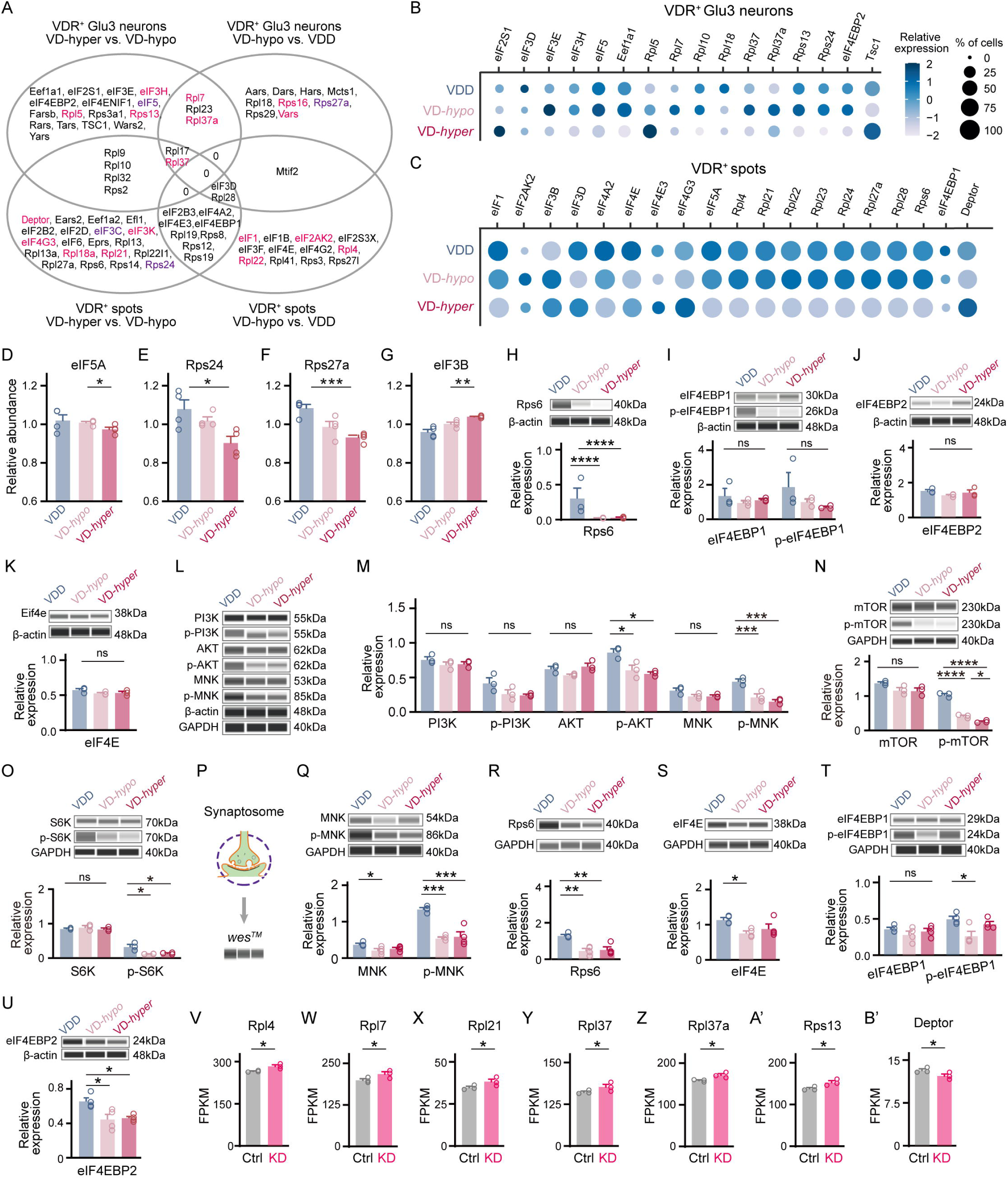
Vitamin D regulates translation via transcriptional control, as well as mTOR signaling in olfactory neurons and synapses. **A.** Venn diagram showing example overlapping DEGs from snRNA-seq (comparing VDR□ Glu3 groups: VD-*hyper* vs. VD-*hypo* and VD-*hypo* vs. VDD) and spatial transcriptomics (comparing VDR□ spots: VD-*hyper* vs. VD-*hypo* and VD-*hypo* vs. VDD). Genes marked in pink represent DEGs confirmed by RNA sequencing in control vs. VDR knockdown (VDR-KD) mice; genes in purple correspond to DEPs from proteomic analysis (VD-*hyper* vs. VD-*hypo* or VD-*hypo* vs. VDD); genes in blue are confirmed by both techniques. **B-C.** Dot plots showing expression of translation-related DEGs in VDR□ Glu3 neurons from snRNA-seq (**B**) and VDR□ spots from spatial transcriptomics (**C**) across three groups (VD-*hyper*, VD-*hypo*, and VDD). Dot size represents the percentage of cells expressing the gene; color indicates averaged scaled expression levels. **D-G.** Relative expression levels of eIF5A (**D**), Rps24 (**E**), Rps27a (**F**), and eIF3B (**G**) in LC-MS-based proteomic analysis (n=4 mice per group; unpaired t-test). **H-K.** Levels of mTOR signaling components, including Rps6 (**H**), eIF4EBP1 and p-eIF4EBP1 (**I**), eIF4EBP2 (**J**), and eIF4E (**K**), in the OB of the three mouse groups, measured by *wes* (n=3 mice per group; one-way ANOVA with Tukey’s multiple comparisons). **L-O.** Levels of regulators of mTOR signaling (**L-M**), mTOR and p-mTOR (**N**), and downstream targets S6K and p-S6K (**O**) in the OB of the three mouse groups, measured by *wes* (n=3 mice per group; one-way ANOVA with Tukey’s multiple comparisons). **P.** Experimental schematic of whole OB synaptosome preparations and protein-level validation using quantitative *wes*. **Q-U.** Components of mTOR signaling, including MNK and p-MNK (**Q**), Rps6 (**R**), eIF4E (**S**), eIF4EBP1 and p-eIF4EBP1 (**T**), and eIF4EBP2 (**U**), in synaptosome preparations from the OB of three mouse groups, measured by *wes* (n=4 mice per group; one-way ANOVA with Tukey’s multiple comparisons). **V-B’.** Relative mRNA expression of Rpl4 (**V**), Rpl7 (**W**), Rpl21 (**X**), Rpl37 (**Y**), Rpl37a (**Z**), Rps13 (**A’**), and Deptor (**B’**) in control (Ctrl) and VDR-KD mice, analyzed by RNA sequencing (n=3 mice per group; statistical analysis conducted using standard pipelines for RNA-seq data, with normalization and multiple testing correction applied). Each symbol represents a biological replicate. Data are presented as mean ± SEM. Significance levels: *P < 0.05, **P < 0.01, ***P < 0.001, ****P < 0.0001; ns indicates no statistical significance. FPKM: Fragments per kilobase million.

Interestingly, many translation initiation factors—including *eIF1*, *eIF3D*, *eIF4A2*, *eIF4E*, and *eIF5A*—were most highly expressed in VDR^+^ Glu3-containing spots from the VDD group, whereas most differentially expressed ribosomal proteins (*Rpl4*, *Rpl21*, *Rpl22*, *Rpl23*, *Rpl24*, *Rpl27a* and *Rpl28*) showed the lowest expression in VDR^+^ Glu3-containing spots from the VD-*hyper* group (Figure 5C). These results suggest that VDD is associated with elevated translation activity, while high-dose VitD supplementation suppresses translation levels. Notably, molecules associated with mTOR signaling—*TSC1*, eIF4E binding protein 1 (*eIF4EBP1*), eIF4E binding protein 2 (*eIF4EBP2*), *Deptor* and *Rps6*—also displayed differential expression in VDR^+^ Glu3 neurons or VDR^+^ Glu3-containing spots upon varying serum VitD levels, implicating mTOR signaling in the VDR-mediated regulation of translation.

Quantitative proteomic analysis of DEPs in the OB of VDD, VD-*hypo* and VD-*hyper* groups using LC-MS revealed that eIF5A, Rps24, and Rps27a exhibited the lowest expression in the VD-*hyper* group (Figure 5D-F, P<0.05 for eIF5A and Rps24, P<0.001 for Rps27a, unpaired t-test). In contrast, eIF3B was significantly upregulated in the VD-*hyper* group (Figure 5G, P<0.01, unpaired t-test). These findings were consistent with results from snRNA-seq and spatial transcriptomics.

Given the differential expression of mTOR signaling components (*eIF4EBP1*, *eIF4EBP2*, eIF4E and *Rps6*) under varying VitD levels, we performed quantitative western blot (*wes*^TM^) analysis of these proteins in the OBs of VDD, VD-*hypo*, and VD-*hyper* mice. Rps6, a downstream effector of mTOR signaling that directly regulates translation, was significantly upregulated in the VDD group compared the VD*-hypo* and VD-*hyper* groups (Figure 5H, P<0.0001, one-way repeated ANOVA with Tukey’s multiple comparisons post hoc test), consistent with enhanced translation activity during VitD deficiency, likely associated with hyperactivation of mTOR signaling. However, eIF4EBP1, phosphorated eIF4EBP1 (p-eIF4EBP1), eIF4EBP2 and eIF4E showed no differential expression across the three groups (Figure 5I-K).

To explore the upstream mechanisms driving mTOR signaling activation in the VDD group, we investigated known upstream regulators^[46]^. We observed a significant increase in phosphorated AKT serine/threonine kinase (p-AKT), phosphorated mitogen-activated protein kinase-interacting kinase (p-MNK), phosphorylated mTOR (p-mTOR) and phosphorylated ribosomal protein S6 kinase (p-S6K) in the VDD group compared to the VitD-supplemented groups (Figures 5L-O, one-way repeated ANOVA with Tukey’s multiple comparisons post hoc test). However, the expression levels of PI3K, phosphorated PI3K (p-PI3K), AKT, MNK, mTOR and S6K levels were unchanged across the three groups (Figures 5L-O), suggesting that mTOR signaling activation under VitD deficiency is associates with the activation of AKT pathways. Collectively, these findings not only validated the earlier sequencing data but also indicated that increased translation during VitD deficiency is, at least in part, mediated by mTOR signaling activation.

We next investigated whether the observed upregulation of translation during VitD deficiency also occurs at the synaptic level. Quantitative analysis of synaptosome proteins revealed a significant increase in MNK, p-MNK, Rps6, eIF4E, and p-eIF4EBP1 in the VDD group (Figures 5P-T, one-way repeated ANOVA with Tukey’s multiple comparisons post hoc test). Moreover, eIF4EBP2 was significantly more highly expressed in the VDD group compared to the VitD-supplemented groups (Figure 5U, P<0.05, one-way repeated ANOVA with Tukey’s multiple comparisons post hoc test), suggesting a more complexed mTOR signaling at the synapse level. These results indicate that mTOR signaling drives an upregulation of local translation at synapses during VitD deficiency, which may directly contribute to changes in synaptic protein composition and functions.

To determine whether the observed translational changes are mediated by the VDR, we performed viral-mediated knockdown (KD) of VDR in the OB using the SaCas9 system. This intervention led to a significant upregulation of transcripts encoding ribosomal proteins, such as *Rpl4*, *Rpl7*, *Rpl21*, *Rpl37*, *Rpl37a*, and *Rps13* (Figures 5V-A’, P<0.05). Interestingly, *Deptor*, a component of the mTOR complex, was significantly decreased in the OBs of VDR-KD mice (Figure 5B’, P<0.05). These findings underscore the critical role of VDR in modulating translation-related processes in a VitD dose-dependent manner, linking its regulatory function to the mTOR signaling pathway.

### VDR Knockdown in the OB impairs olfactory function in Mice

So far, we have demonstrated that VitD regulates the expression of molecules critically involved in the synaptic and translational processes in the OB in a VDR-dependent manner. Given that mice with VitD deficiency exhibit impaired olfactory function, we next investigated whether VDR knockdown could reproduce similar behavioral deficits. To address this, we performed a series of behavioral experiments on mice with viral-mediated VDR knockdown in the OB (Figure 6A). Both the control and VDR-KD groups had similar body weights and showed comparable performance in the open field test, including total distance traveled, time spent in the center, and travel speed (Figure 6B-E). These results suggest that neither the viral injection procedure nor VDR knockdown in the OB affects general motor function.

**Figure 6.**
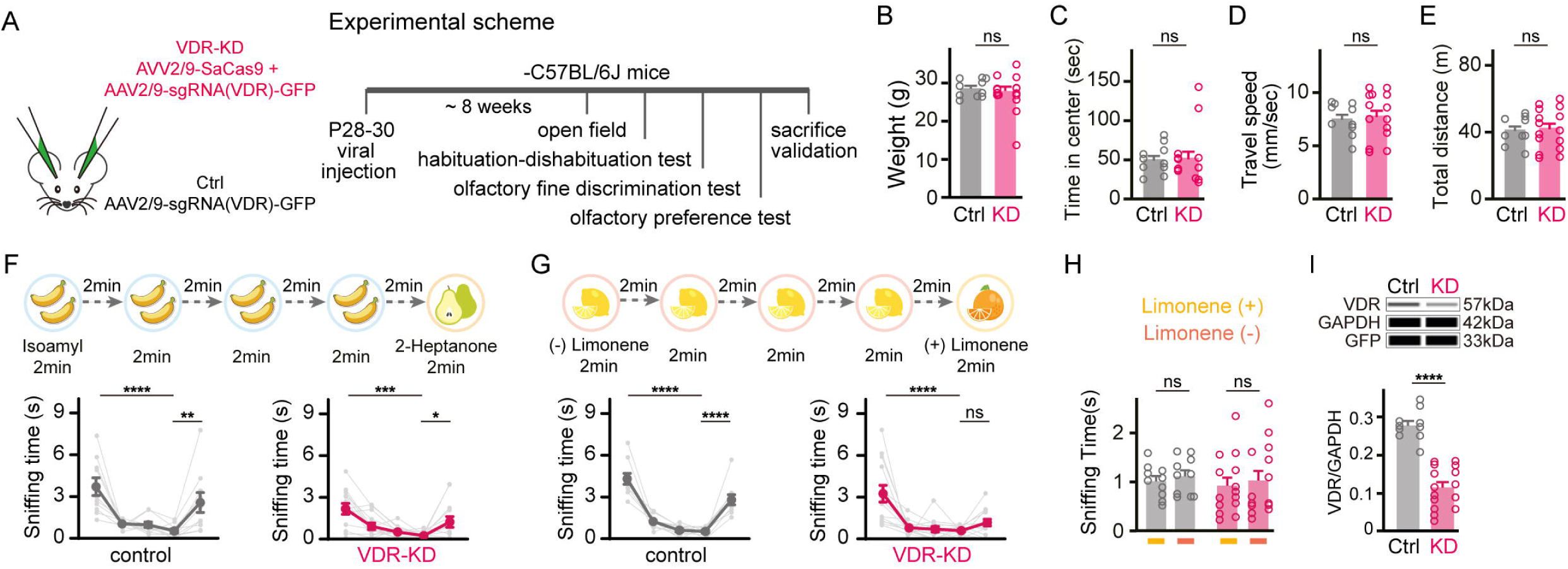
VDR knockdown in the OB impairs olfactory function in mice. **A.** Experimental schematic of CRISPR-Cas9-mediated VDR-KD in the mouse OB and subsequent behavioral tests. **B-E.** Body weight (**B**) and measurements from the open field test, including time to enter the center area (**C**), average movement speed (**D**), and total movement distance (**E**), in VDR-KD and control groups (unpaired t-test). **F-G.** Olfactory habituation-dishabituation tests: sniffing time of VDR-KD and control mice in response to odor pairs, including isoamyl acetate vs. 2-heptanone (**F**) and (L)-limonene vs. (D)-limonene (**G**) (one-way ANOVA with Tukey’s correction for multiple comparisons). **H.** Sniffing time for (L)-limonene and (D)-limonene in the olfactory preference test for VDR-KD and control groups (unpaired t-test). **I.** The *wes* image and quantification of VDR expression in VDR-KD and control (Ctrl) groups, measured after completion of behavioral tests (unpaired t-test). Each symbol represents a biological replicate (n=10 mice per group). Data are presented as mean ± SEM. Significance levels: *P < 0.05, **P < 0.01, ***P < 0.001, ****P < 0.0001; ns indicates no statistical significance.

In the olfactory habituation test, both groups exhibited a significant decrease in sniffing time between the first and the fourth presentations of isoamyl acetate (Figure 6F, P<0.0001 for control; P<0.001 for VDR-KD, unpaired t-test) and a significant increase in sniffing time when the odor was switched to 2-heptanone in the fifth trial (Figure 6F, P<0.01 for control; P<0.05 for VDR-KD, unpaired t-test). However, when presented with two highly similar odors (L)-limonene and (D)-limonene, the control group showed significantly increased in sniffing time during the fifth trial when the odor changed from (L)-limonene to (D)-limonene (Figure 6G, P<0.0001, unpaired t-test). In contrast, the VDR-KD group displayed no change in sniffing time between the fourth and fifth trials (Figure 6G, unpaired t-test), indicating impaired fine odor discrimination. Notably, both groups showed similar preference for (L) - and (D)-limonene, confirming that the mice had no intrinsic preference for either odor (Figure 6H, unpaired t-test). Finally, the success of VDR knockdown was validated by quantitative western blot (Figure 6I, P<0.0001, unpaired t-test). Collectively, these findings suggest that reduced VDR expression in the OB impairs olfactory function in mice. This impairment may be linked to the disruption of VDR-mediated regulation of synaptic and translational pathways, highlighting the critical role of VitD signaling in maintaining olfactory circuit integrity and function.

### mTOR inhibition rescues olfactory deficits in VitD-deficient mice by normalizing translation and excitatory synaptic protein expression

Given that VDD mice exhibited impaired olfactory function alongside increased translation and synaptic protein expression, and considering the association of the mTOR signaling pathway with the VitD-dependent translational regulation, we next investigated whether suppressing overactivated translation through mTOR inhibition could rescue synaptic and behavioral deficits caused by VitD deficiency. To this end, we treated VDD mice with rapamycin, a specific mTOR inhibitor, and conducted a series of experiments (Figure 7A).

**Figure 7.**
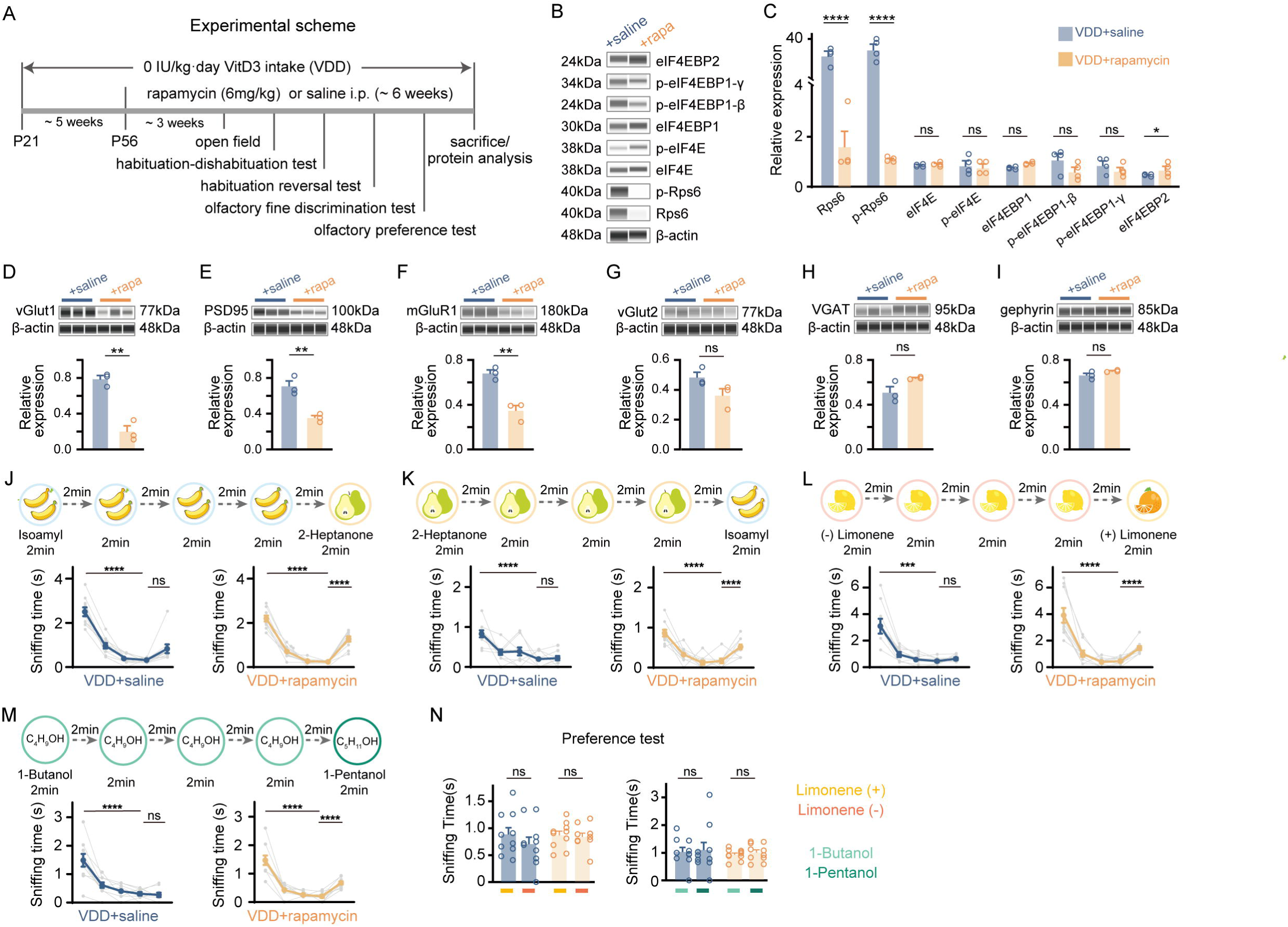
mTOR inhibition rescues olfactory deficits in vitamin D-deficient mice by normalizing translation and excitatory synaptic protein expression. **A.** Experimental scheme of behavioral tests conducted on rapamycin-treated (rapa) and control (saline) groups. VDD mice were treated with 6 mg/kg rapamycin or saline three times per week starting from P56 for six weeks, while behavioral tests started three weeks after rapamycin treatment. **B-C.** Protein expression levels of mTOR-associated translational machinery components in the OB of rapamycin-treated and control groups, measured by *wes* (n=4 mice per group; unpaired t-test). **D-I.** Expression levels of synaptic proteins, including vGlut1 (**D**), PSD95 (**E**), mGluR1 (**F**), vGlut2 (**G**), VGAT (**H**), and gephyrin (**I**), in synaptosome preparations from OBs of the two groups, measured by *wes* (n=3 mice per group; unpaired t-test). **J-M.** Sniffing time in the olfactory habituation-dishabituation test (**J, L, M**) and olfactory habituation reversal test (**K**) (n=10 mice per group; one-way ANOVA with Tukey’s correction for multiple comparisons). Odor pairs included isoamyl acetate vs. 2-heptanone (**J, K**), (L)-limonene vs. (D)-limonene (**L**), and 1-butanol vs. 1-pentanol (**M**). **N.** Sniffing times for (L)-limonene vs. (D)-limonene (left) and 1-butanol vs. 1-pentanol (right) in the olfactory preference test (n=10 mice per group; unpaired t-test). Each symbol represents a biological replicate. Data are presented as mean ± SEM. Significance levels: *P < 0.05, **P < 0.01, ***P < 0.001, ****P < 0.0001; ns indicates no statistical significance.

First, rapamycin treatment significantly reduced the protein levels of Rps6 and increased eIF4EBP2 levels in the OB of VDD mice, without altering the expression of eIF4E, eIF4EBP1, or p-EIF4EBP1 (Figures 7B-C, P<0.05 for Eif4ebp2; P<0.0001 for Rps6 and P-Rps6, unpaired t-test). These results indicate that rapamycin effectively downregulates mTOR-associated translational machinery. Furthermore, in synaptosome fractions, the expression of excitatory synaptic markers—vGlut1, mGluR1, and PSD95—was significantly reduced following rapamycin treatment (Figures 7D-F, P<0.01, unpaired t-test), while vGlut2 and markers of inhibitory synapse (VGAT and gephyrin) remained unchanged (Figures 7G-I). These findings suggest that mTOR-dependent translation plays a specific role in modulating the expression of excitatory synaptic proteins under conditions of VitD deficiency.

Next, we examined whether rapamycin-induced translational downregulation and the associated reduction in excitatory synaptic proteins could rescue olfactory deficits in VDD mice. Although rapamycin treatment led to a decrease in body weight (Figure S9A, P<0.01, unpaired t-test), the brain-to-body weight ratio remained unaffected (Figure S9B). Additionally, general motor function, as measured by the travel distance and speed in the open field, were unaffected; however, rapamycin-treated mice spent significant less time in the center zone (Figures S9C-S9E, P<0.05 for time in the center zone, unpaired t-test). In the olfactory habituation test, both groups showed a significant decrease in sniffing time between the first and the fourth presentations of isoamyl acetate (Figure 7J, P<0.0001, unpaired t-test). However, only the rapamycin-treated VDD group exhibited a significant increase in sniffing time upon odor switch to 2-heptanone in the fifth trial (Figure 7J, P<0.0001, unpaired t-test). Similar results were observed when the odor sequence was reversed (Figure 7K, P<0.0001, unpaired t-test). Moreover, the rapamycin-treated VDD group demonstrated improved discrimination of structurally similar odor pairs, such as (L)-limonene and (D)-limonene, as well as 1-butanol and 1-pentanol (Figures 7L-M, P<0.0001, unpaired t-test), indicating enhanced olfactory function compared to the saline-treated group. Importantly, both groups exhibited similar intrinsic preferences for the odor pairs, confirming that rapamycin treatment did not alter odor preference biases (Figure 7N). Taken together, these findings suggest that mTOR inhibition via rapamycin rescues olfactory deficits in VDD mice by normalizing translation and excitatory synaptic protein expression, further supporting the role of mTOR signaling in mediating the effects of VitD deficiency on synaptic and behavioral function.

## Discussion

Our study uncovers a mechanistic link between VitD signaling and olfactory function through mTOR-dependent synaptic remodeling. We found that the VDR is enriched in the OB, particularly in tufted cells of the GL and EPL, with expression levels increasing during postnatal development. Using dietary manipulation models, we demonstrated that VitD bidirectionally modulates olfactory function—deficiency impaired odor discrimination, while supplementation enhanced sensitivity. Transcriptomic and proteomic analyses revealed that VDR regulates synaptic protein expression in olfactory neurons through both transcriptional control and mTOR-mediated translational mechanisms. VitD deficiency drove excessive upregulation of excitatory synaptic components (e.g., vGlut1, PSD95), a phenotype partially reversible by mTOR inhibition with rapamycin. These findings position VitD as a key regulator of neural circuit refinement through translational control, extending its classical endocrine roles.

The connection between VitD and olfactory dysfunction is clinically relevant, given that olfactory deficits are early markers of certain types of neurodevelopmental and neurodegenerative disorders^[47,48]^. The mechanistic overlap with autism spectrum disorder (ASD) is particularly striking, as high-confidence ASD risk genes (*FMR1*, *TSC1/2*, *PTEN*, *CYFIP1*, etc.) converge on mTOR-dependent translational pathways^[49,50]^. VitD deficiency recapitulated core features of ASD synaptic pathology, including elevated excitatory protein synthesis similar to *FMR1* knockout models and *TSC1*-associated mTOR hyperactivity^[51–54]^. Our demonstration that VitD deficiency exacerbates mTOR-driven synaptic protein translation mirrors core aspects of these ASD models, providing a mechanistic framework for understanding how VitD status might influence neurodevelopmental trajectories. While clinical trials of VitD supplementation in ASD have yielded variable results^[12,13]^, our data suggest that patient stratification based on mTOR pathway activation status could refine therapeutic targeting.

At the molecular level, VitD modulates protein expression through transcriptional control, as well as the PI3K/AKT/mTOR/translation axis. VitD deficiency not only increased the transcript levels of specific synaptic proteins, but also elevated p-AKT and p-MNK levels, suggesting ERK-dependent translational activation downstream of enhanced excitatory transmission^[26,50]^. Notably, we observed selective synaptic enrichment of eIF4E in VitD-deficient mice, indicating localized dysregulation of protein synthesis machinery^[54]^. These findings align with the observed upregulation of excitatory synaptic proteins in VitD deficiency, supporting a novel role for VitD in synaptic translational control. Beyond mTOR-mediated effects, VitD may directly regulate translation by influencing the transcription of translation-related factors. Our large-scale transcriptomic analysis revealed differential expression of several translation regulators— including eIF2B2, eIF4G2, eIF5A, Rpl22, Rpl37a, Rps3, and Rps6K (S6K)—many of which possess vitamin D response elements (VDREs)^[15]^. This suggests potential transcriptional regulation of translation factors by VitD. However, further investigations should aim to precisely delineate how VitD status modulates the translational process, whether through transcriptional regulation of pathway components, and/or through coordinating activity-dependent translational responses. Moreover, the spatiotemporal dynamics of translation machinery trafficking, particularly the mechanisms underlying its synaptic enrichment under conditions of VitD depletion, require systematic characterization.

Using molecular profiling, we identified VitD regulates signaling pathways essential for excitatory-inhibitory balance within olfactory bulb microcircuits centered around the tufted cells. Importantly, VitD deficiency induced a marked imbalance in synaptic protein composition, characterized by disproportionate elevation of excitatory markers (vGlut1, PSD95) relative to inhibitory components. This dysregulation likely perturbs the precisely-timed network oscillations (gamma/theta coupling) and temporal spike patterns known to encode odor identity and discrimination^[55–57]^. Notably, the synaptic protein expression changes identified in the olfactory bulb diverged from those reported in hippocampal or cortical circuits—whether studied under conditions of gestational VitD deficiency^[58,59]^ or adult VitD supplementation^[21,22]^. This region- and cell-type-specificity further underscores that VitD’s synaptic regulation is spatiotemporally constrained. Key unresolved questions include how VitD-dependent modulation varies across circuits, whether these changes alter tufted-cell excitability and synaptic transmission, and if excitation/inhitibition imbalance compromises population-level odor coding. Resolving these mechanisms—though beyond this study’s scope—is vital for understanding how nutritional state, via VitD, impacts synaptic homeostasis and sensory processing.

Collectively, our findings establish a functionally specialized subpopulation of tufted cells as a critical VitD-sensitive node in olfactory circuit regulation, wherein mTOR-dependent control of synaptic protein translation serves as a key mechanistic link between nutritional state and sensory processing. Beyond advancing fundamental knowledge of nutrient-sensitive synaptic regulations, these insights highlight novel therapeutic opportunities—whether through dietary optimization, VitD supplementation, or strategic mTOR modulation—as potential stand-alone or adjunctive interventions in neurodevelopmental and neurodegenerative disorders marked by synaptic dysfunction.

## Resource availability

The structured datasets—including snRNA-seq, spatial transcriptomics, bulk RNA sequencing— are publicly available through the NCBI repository (BioProjectID : PRJNA1248507). The mass spectrometry proteomics data have been deposited to the ProteomeXchange Consortium (https://proteomecentral.proteomexchange.org) via the iProX partner repository with the dataset identifier PXD062768. Additional raw datasets not deposited in NCBI can be obtained upon request from the corresponding author or accessed via the Mendeley Data repository (https://data.mendeley.com/preview/mfs9283rt9?a=883de594-374f-4911-82c3-2ce3f4efd2e2).

## Ethics approval

All animal experiments were performed according to the Ethics Committee of Hainan Medical University (HYLL-2022-259 and HYLL-2024-173) and conformed to the ethical principles of welfare of laboratory animals.

## Consent to participate

This study didn’t involve human subjects.

## Consent for publication

This study didn’t involve patients.

## Authors’ contributions

L.X. and W.X. conceived and designed the study. L.X., R.P.C., C.R.H., and Y.X.S. wrote the manuscript. R.P.C. and C.R.H. jointly performed multi-omics data analysis and conducted quantitative *wes* validation experiments. R.P.C. carried out olfactory behavior tests and *VDR* knockdown experiments, with assistance from B.Q.S., and performed rapamycin treatment studies. Y.X.S. executed FISH and immunohistochemistry assays, along with snRNA-seq and spatial transcriptomics experiments. P.W.B. established the dietary vitamin D mouse model and performed bulk RNA sequencing and proteomics analyses. H.M.H. contributed to data analysis and figure preparation. Y.D. and Z.Q.L. participated in establishing the vitamin D dietary mouse model and provided technical assistance. All authors critically reviewed and approved the final manuscript.

## Acknowledgements

We thank Genergy Biotechnology (Shanghai) Co., Ltd., for assisting the snRNA-seq and spatial transcriptomics sequencing analysis. We thank Wang Meijuan, Li Hongai, and Lin Xinmei for technique assistant and scientific discussions. We thank Dr. Chen Jingke, Dr. Han Yunyun and Dr. Feng Shi for kind suggestions to our manuscript.

## Declaration of generative AI and AI-assisted technologies in the writing process

During the preparation of this work the authors used deepseek-ai/DeepSeek-V3 in order to improve the language and readability. After using this tool/service, the authors reviewed and edited the content as needed and take full responsibility for the content of the publication.

## STAR Methods

### Mice

Three-week-old male C57BL/6J mice were obtained from Hunan SJA Laboratory Animal Co., Ltd. Vglut2-ires-Cre (Slc17a6^tm2(cre)Lowl^/J, Jacksonlab Strain #016963) transgenic mice^[36]^ were generously provided by Dr. Xu Chun’s research group at the Institute of Neuroscience, Chinese Academy of Sciences, Shanghai. Mice were housed in individually ventilated cages under a 12-hour light/12-hour dark cycle, with five mice per cage, and provided *ad libitum* access to water. Environmental conditions were maintained at a temperature of 22 ± 2°C and a relative humidity of 55 ± 10%. All experimental procedures were conducted in compliance with the ethical guidelines approved by the Ethics Committee of Hainan Medical University (Approval Numbers: HYLL-2022-259 and HYLL-2024-173). Unless otherwise specified, mice were fed the standard AIN-93G diet *ad libitum*.

### Vitamin D3 supplementation

Mice with gradient vitamin D3 supplementation were established as described before. In brief, mice were weaned at P21 and fed customized diets formulated to provide daily vitamin D□ intakes of 0, 500, or 2000 IU/kg for a minimum of 12 weeks, corresponding to the VDD group, VD-*hypo* group and VD-*hyper* group. Six custom diets, based on the standard AIN-93G formulation with vitamin D□ concentrations of 0, 1.00×10³, 2.50×10³, 5.00×10³, 1.00×10□, and 2.00×10□ IU/kg, were obtained from Jiangsu Xietong Bioengineering Co., Ltd. (Nanjing, China).

### Rapamycin administration

Starting at P56, rapamycin (R-5000, LC Laboratories, USA) was administered to VDD mice via intraperitoneal injection. A 20 mg/mL stock solution of rapamycin was prepared in dimethyl sulfoxide (DMSO, 60313ES60, Yeasen, China) and stored at −80°C. Prior to injection, the stock solution was diluted to 0.6 mg/mL using sterile 0.9% saline (BL158A, Biosharp, China). Mice received intraperitoneal injections of rapamycin at 6 mg/kg, three times per week, for six weeks. Control mice received vehicle (DMSO in saline) on the same schedule.

### Measurement of the serum 25(OH)D

At ∼14 weeks of age, the mice were anesthetized using isoflurane, and approximately 500µL of blood was collected via orbital enucleation. Serum levels of 25-hydroxyvitamin D_2_ [25(OH)D_2_] and 25-hydroxyvitamin D_3_ [25(OH)D_3_) were quantified using LC-MS (API 3200 LC-MS, AB Sciex) at Daan Gene Company Limited (Guangzhou, China). Total serum 25(OH)D levels were calculated as the sum of 25(OH)D_2_ and 25(OH)D_3_ concentrations.

### Stereotactic injections

Four-week-old Mice were anesthetized with 2-3% isoflurane and placed in a stereotactic frame (Reward Co.). A total of 200∼300 nl of adeno-associated virus (AAV) was bilaterally injected into the olfactory bulbs using the following coordinates: medio-lateral (M/L) ± 0.6 mm, anterior-posterior (A/P) +4.5 mm, and dorso-ventral (D/V) −3.0 mm. AAV2-hSYN-DIO-EGFP-KASH-WPRES (titer: 2.75× 10^12^ vg/mL) was a gift from Dr. Peter Scheiffele (Biozentrum, Basel, Switzerland). Two weeks post-injection, mice were processed for immunohistochemical analysis. For VDR knockdown experiments, AAV2/9-U6-sgRNA(VDR)-EF1α-EGFP-WPRE-pA (titer: 2.09×10^12^ vg/mL) and AAV2/9-hSyn-saCas9-3HA-pA (titer: 5.52×10^12^ vg/mL) were purchased from Brain VTA Technology Company Limited (Wuhan, China). Behavioural assays were conducted 8 weeks post-injection, and a separate cohort for sequencing experiments were performed 4□weeks post-injection.

### Nissl Staining

Mice were anesthetized with isoflurane and transcardially perfused with 4% paraformaldehyde (PFA). After dehydrations, OBs were sectioned at 50μm thickness using a microtome (Leica CM1950, Wetzlar, Germany). Sections were stained with cresyl violet (Nissl) staining solution (C0117, Beyotime, China), dehydrated through a graded ethanol series, cleared in xylene, and mounted. Images were acquired using a light microscope (Murzider, Z530) and analyzed with Fiji software.

### Immunohistochemistry

Brain sections were blocked for 1 hour at room temperature in Tris-buffered saline (TBS) containing 0.3% Triton X-100 (30188928 Sinopharm Chemical Reagent Co., China) and 10% goat serum (Zhongshan Jinqiao Biotechnology, Beijing, China). Sections were then incubated overnight at 4°C with primary antibodies diluted in TBS containing 0.3% Triton X-100 and 5% goat serum. After washing, sections were incubated with secondary antibodies at room temperature for 1 hour. Nuclei were counterstained with DAPI (BL105A, Biosharp, China). Images were acquired using a confocal microscope (TCS SP8, Leica, Germany). The following primary antibodies were used: rabbit anti-vitamin D receptor (1:500; Cell Signaling Technology, 12550S); rabbit anti-GFP (1:10000; Invitrogen, A11122); rabbit anti-calbindin D-28k (calb2) (1:5000; Swant CB38); mouse anti-calretinin (calb1) (1:5000; 6B3, Swant); guinea pig anti-somatostatin-28 (1:500; Synaptic Systems, 366004); mouse-anti-parvalbumin (1:5000; Swant 235); rabbit anti-EGFP (1:1000; Invitrogen A-11122); mouse-anti-tyrosine hydroxylase (1:500; Sigma MAB318); guinea pig anti-S100b (1:200; Synaptic Systems, 287004). Secondary antibodies used were: goat anti-rabbit Alexa Fluor 568 (1:500; Invitrogen, A11011); goat anti-mouse Alexa Fluor 488 (1:500; Invitrogen, A11029); goat anti-chicken Alexa Fluor 633 (1:500; Invitrogen, A11039).

### FISH

Adult mice were anesthetized and decapitated, and brains were rapidly extracted and sectioned at 20μm thickness using a frozen microtome (Leica CM1950UV, Leica, Germany). RNAscope® in situ hybridization was performed according to the manufacturer’s protocol (323100, Advance Cell Diagnostics (ACD), Hayward, CA) and previously described method. Briefly, sections were pretreated and incubated for 30 minutes on HybEz™ Hybridization System (321720, ACD Hayward, CA). Sections were then hybridized with a VDR probe (C1#524511) in a HybEZ oven (ACD) at 40°C for 2 hours. Negative (320871, ACD) and positive (320881, ACD) control probes were used under the same conditions. Nuclei were counterstained with DAPI (BL105A, Biosharp, China). Images were acquired using a confocal microscope (TCS SP8, Leica, Germany) and analyzed using Fiji software. For quantification, two olfactory bulb sections per mouse (n=3 mice, 6 sections total) were analyzed, with 5–6 microscopic fields captured per section (30–36 images per experiment). DAPI-positive cells were counted in each anatomical layer (GL, EPL, MCL, and GCL). Cells were considered positive for VDR mRNA expression if they exhibited more than five probe signal spots localized within the nucleus or perinuclear compartment (signal within 2–3 µm²).

### RNA extraction and real-time quantitative reverse transcription PCR (RT-qPCR)

Total RNA was purified using Eastep® Super Total RNA Extraction Kit (Promega LS1040, Shanghai, China). Reverse transcription was performed using the High-Capacity cDNA Reverse Transcription Kit (Applied Biosystems 4368814, Foster City, CA, USA) following the manufacturer’s instructions. Quantitative PCR (qPCR) was conducted on a CFX96 real-time detection system (Bio-Rad, Hercules, CA, USA) using PowerUp™ SYBR™ Green Master Mix (Applied Biosystems A25742, Carlsbad, CA, USA). Primer specificity was confirmed by melting curve analysis and agarose gel electrophoresis. Relative gene expression was calculated using the 2^−ΔΔCT^ method, normalized to GAPDH. All measurements were performed in triplicate. Primer sequences: VDR: Forward, TCAAACTCTGATCTGTACACCC, Reverse, TGGATGCTGTAACTGACAAGAT; GAPDH: Forward, GGTTGTCTCCTGCGACTTCA; Reverse, TGGTCCAGGGTTTCTTACTCC.

### Protein extraction and synaptosome preparation

Brain tissues were homogenized in RIPA Lysis Buffer (20101ES60, Yeasen, China) supplemented with phenylmethylsulfonyl fluoride (BL507A, Biosharp, China) and a phosphate inhibitor cocktail (BL615A, Biosharp, China). Homogenates were centrifuged at 14,000 rpm for 10 minutes and the supernatant was collected for protein quantification using the BCA™ Protein Assay (20201ES76, Yeasen, China).

Synaptosome fraction were isolated using the Syn-PER Synaptic Protein Extraction Reagent (Thermo Fisher Scientific, Waltham, MA). Tissues were homogenized in 100□µl of Syn-PER reagent and 1□µl of Halt Protease and phosphatase inhibitor cocktail (Thermo Fisher Scientific). Homogenates were centrifuged at 500×g for 10□min at 4□°C, and the supernatant was further centrifuged at 15,000×g for 20□min at 4□°C. The resulting synaptosome pellet was resuspended in 40 µl of Syn-PER reagent with Halt inhibitor cocktail.

### *Wes*^TM^ Simple Western assays (Protein Simple *Wes*)

Capillary Western blot analysis was performed using the Jess Simple Western system (Bio-Techne, Minneapolis, MN, USA) as previously described. Capillary cartridges (Protein Simple, SM-W004-1), anti-rabbit detection module chemiluminescence (Protein Simple DM-001), anti-mouse detection module chemiluminescence (Protein Simple DM-002) and EZ standard pack (Protein Simple PS-ST01EZ-8) were used according to the manufacturer’s instructions. The following antibodies were used: mouse anti-β-actin (1:200, Novus, NB600-501), rabbit anti-AKT (1:200, Cell Signaling Technology, 4691), rabbit anti-p-AKT (1:20, Cell Signaling Technology, 4060), rabbit anti-eIF4E (1:100, Cell Signaling Technology, 2067), rabbit anti-eIF4EBP1 (1:100, Cell Signaling Technology, 9644), rabbit anti-p-eIF4EBP2 (1:200, Cell Signaling Technology, 2855), Rabbit anti-eIF4EBP2 (1:20, CST, 2845), rabbit anti-GAPDH (1:5000, Cell Signaling Technology, 5174), mouse anti-gephyrin (1:4000, Synaptic Systems, 147111), rabbit anti-mGluR1 (1:500, Cell Signaling Technology, 12551), rabbit anti-MNK (1:100, Cell Signaling Technology, 2195), rabbit anti-p-MNK (1:20, Cell Signaling Technology, 2111), mouse anti-mTOR (1:1000, CST, 4517), rabbit anti-p-mTOR (1:50, CST, 5536), rabbit anti-PI3K (1:3000, Cell Signaling Technology, 4249), rabbit anti-p-PI3K (1:20, Cell Signaling Technology, 17366), rabbit anti-PSD95 (1:100, Cell Signaling Technology, 3409), rabbit anti-Rps6 (1:100, Cell Signaling Technology, 2217), rabbit anti-S6K (1:200, CST, 33475), rabbit anti-p-S6K (1:50, CST, 9234), mouse anti-synapsin1 (1:7000, Synaptic Systems, 106011), rabbit anti-VDR (1:500, Cell Signaling Technology, 12550), mouse anti-VGAT (1:20, Synaptic Systems, 131011), mouse anti-vGlut1 (1:7000, Synaptic Systems, 135011), rabbit anti-vGlut2 (1:100, Cell Signaling Technology, 16066).

### RNA library construction and sequencing (Bulk RNA-sequencing)

Total RNA of olfactory bulbs was isolated using the RNeasy Lipid Tissue Mini Kit (Qiagen, Hilden, Germany). RNA quality was assessed by gel electrophoresis and Qubit (Thermo, Waltham, MA, USA). Libraries were prepared using the VAHTS Stranded mRNA-seq Library Prep Kit for Illumina (Vazyme, China) and sequenced on the Illumina Novaseq 6000 platform. Raw data were processed using Skewer, and quality was assessed with FastQC (v0.11.2). Clean reads were aligned to the reference genome using STAR (2.5.3a). Transcript expression was quantified as FPKM (Fragments Per Kilobase of exon model per Million mapped reads) using StringTie (v1.3.1c). Differentially expressed genes (DEGs) were identified using DEGseq2 (v1.16.1) with a false discovery rate (FDR) < 0.01 and fold change ≥2.

### snRNA-seq

Olfactory bulbs from six 15-week-old mice (two per dietary group: VDD, VD-*hypo*, VD-*hyper*) were dissected and processed according to the 10X Genomics Chromium protocol. Nuclei were extracted from mouse olfactory bulb tissue using a protocol adapted from the 10X Genomics Demonstrated Protocol CG00055 (RevA) for sample preparation. Nuclei were loaded onto the 10X Chromium Single Cell Platform at a concentration of 1,000 nuclei/μL, following the literature^[60,61]^ and the manufacturer’s instructions for the Single Cell 3′ library and Gel Bead Kit v.3. Gel beads in emulsion (GEMs) were generated, and barcoding, GEM-RT clean-up, cDNA amplification, and library construction were performed following the manufacturer’s protocol. Libraries were quantified using Qubit and sequenced on the Illumina NovaSeq 6000 instrument with 150-base-pair paired-end reads.

### Unsupervised clustering and visualization

Unsupervised clustering was performed using Seurat package (v2.2) in R. Genes expressed in fewer than two cells were excluded. Cells with >200 genes and <10% mitochondrial genes were retained. Dimensionality reduction was performed using principal component analysis (PCA) based on the top 2,000 most variable genes. A k-nearest neighbor graph was constructed, and clusters were identified using the Louvain Modularity optimization algorithm. Results were visualized using uniform manifold approximation and projection (UMAP) plots. Cells expressing hemoglobin genes were excluded.

### Marker gene identification and cell-type annotation

Differentially expressed genes for each cluster were identified using Seurat’s FindMarkers function with the “bimod” test. Genes with a log2 average expression difference > 0.585 and *P* < 0.05 were considered marker genes. Clusters were annotated using canonical cell-type markers.

### CellphoneDB

Cell-cell communication analysis was performed using CellphoneDB (v5.0.0)^[45]^ for Cluster-3 VDR^+^ Glu3, Cluster-3 VDR-Glu3, and 20 other clusters. Significant interactions (*P* < 0.05) were identified based on normalized cell matrices from Seurat.

### Spatial Transcriptomics analysis

Olfactory bulb sections (10 µm) from 15-week-old mice (one per dietary group: VDD, VD-*hypo*, VD-*hyper*) were processed using the 10X Genomics Visium platform^[62,63]^. These processes included H&E staining, infiltration, and library construction according to the protocol provided by the manufacturer. Libraries were sequenced on the Illumina NovaSeq 6000 platform. The Space Ranger pipeline used to process Visium spatial gene expression data followed the guidelines provided by the manufacturer, including demultiplexing, mm10 mouse reference genome alignment, tissue detection, reference detection, and barcode/UMI counting. High-quality spots (≥200 genes) were analyzed using Seurat’s SCTransform for normalization, PCA for dimensionality reduction, and t-SNE/UMAP for visualization.

### GO analyses

Gene ontology (GO) enrichment analyses were performed using the “clusterProfiler” and “TopGO” R package. P-values were calculated using Fisher’s exact test and adjusted using the Benjamini-Hochberg method, with significance defined as P < 0.05.

### GSEA

GSEA was performed using the following workflow: Raw read counts were first normalized to generate a standardized gene expression matrix, which was then imported into the GSEA platform (v4.3.2, https://www.gsea-msigdb.org/gsea/index.jsp) for enrichment analysis. The method calculates an Enrichment Score (ES) for each predefined pathway based on the ranked distribution of genes across the dataset. Statistical significance of the ES was assessed using a permutation test. The ES values were subsequently normalized across pathways to generate Normalized Enrichment Scores (NES), and significantly enriched pathways were identified after adjusting for multiple testing corrections.

### RCTD

Deconvolution analysis was performed using the RCTD package (v2.0.0)^[44]^. Single-cell RNA-seq data were used to define cell-type-specific gene sets, and spatial transcriptomics data were modeled as a Poisson distribution. The expression values for known cell types in the single-cell data were normalized to their average expression values and summed according to unknown proportional weights, while accounting for the batch random effects from both single-cell and spatial platforms. A hierarchical statistical model was constructed to estimate the proportions of known cell types and to provide the maximum-probability estimates for both single and double cell types.

### Quantitative proteomic analysis by LC-MS

Quantitative 4D label-free whole protein analysis was conducted by Jingjie PTM BioLab (Hangzhou, China). Tissue samples were lysed in buffer containing 8M urea and 1% protease inhibitor cocktail, followed by sonication on ice (3 cycles). Lysates were centrifuged at 12,000 × g for 10 minutes at 4°C to remove debris, and the supernatant was collected for protein quantification using a BCA kit. For trypsin digestion, proteins were reduced with 5 mM dithiothreitol at 56°C for 30 minutes and alkylated with 11 mM iodoacetamide at room temperature for 15 minutes in the dark. Peptide precursor ions and their fragments were analyzed using a Bruker timsTOF Pro mass spectrometer (100–1700 m/z) with Parallel Accumulated Serial Fragmentation (PASEF).

Secondary mass spectrometry data were processed using MaxQuant (v1.6.15.0). The *Mus musculus* reference proteome was used for database searching (Mus_musculus_10090_GCF_000001635.26_GRCm38.p6_20220323.Fasta; 84,985 sequences). A decoy database was included to calculate the false discovery rate (FDR), and a common contamination library was incorporated to account for potential contaminants.

### Open Field Test

Mice were placed in a 50 × 50 × 50 cm chamber for 10 minutes, and their activity was recorded using the VisuTrack animal acquisition and analysis platform (Shanghai Xinruan). The central area was defined as the central quarter of the chamber floor, with the remaining area designated as the edge. Movement traces were analyzed using SuperMaze software (v3.0, Shanghai Xinruan Information Technology Co., Ltd.). The experimental setup was enclosed with curtains, and infrared lighting-maintained illumination at 15 lux.

### Olfactory Habituation-Dishabituation Test and Fine Discrimination Test

Olfactory tests were conducted in a 30 × 20 × 15 cm cage containing approximately 0.5 cm of irradiated woodchip bedding. For the habituation-dishabituation test, mice were acclimated to the cage for 30 minutes. A sterile cotton swab soaked in 0.2% isoamyl acetate (Solarbio, M8040; Xilong Scientific, 123-92-2) diluted in mineral oil was presented for 2 minutes, followed by a 2-minute rest period. This procedure was repeated four times (habituation phase). In the fifth trial, a swab soaked in 0.2% 2-heptanone (Sigma-Aldrich, H810890) was presented. For the reversal test, 0.2% 2-heptanone was presented in four trials, followed by 0.2% isoamyl acetate in the fifth trial. Sniffing time, defined as the duration the mouse’s nose was within 2 cm of the swab, was recorded manually.

For the fine discrimination test, 0.2% (L)-limonene (Sigma-Aldrich, 218367) or 1-butanol (Westlink Science, 71-36-3) was presented in the first four trials, followed by 0.2% (D)-limonene (Sigma-Aldrich, 183164) or 1-pentanol (Westlink Science, 71-41-0) in the fifth trial.

### Olfactory Preference Test

Mice were acclimated to the test cage for 30 minutes. Odor pairs [(L)-limonene and (D)-limonene; 1-butanol and 1-pentanol] were presented simultaneously via cotton swabs for 5 minutes. The time spent sniffing each odor was recorded. The order of odor presentation was randomized for each test.

### Statistical Analysis of Data

Statistical analyses were performed using GraphPad Prism (v10.0.2). Normality was assessed using the Kolmogorov-Smirnov or Shapiro-Wilk tests. Data are presented as mean ± SEM if otherwise stated. Comparisons between two groups were analyzed using an independent sample *t*-test (normal distribution) or Mann-Whitney U test (non-normal distribution). For multiple groups, one-way ANOVA followed by Tukey’s post hoc test was used. Each data point represents a single sample or animal. Statistical significance was set at *P* < 0.05, with *P* < 0.05, 0.01, 0.001, and 0.0001 denoted by *, **, ***, and ****, respectively.

---

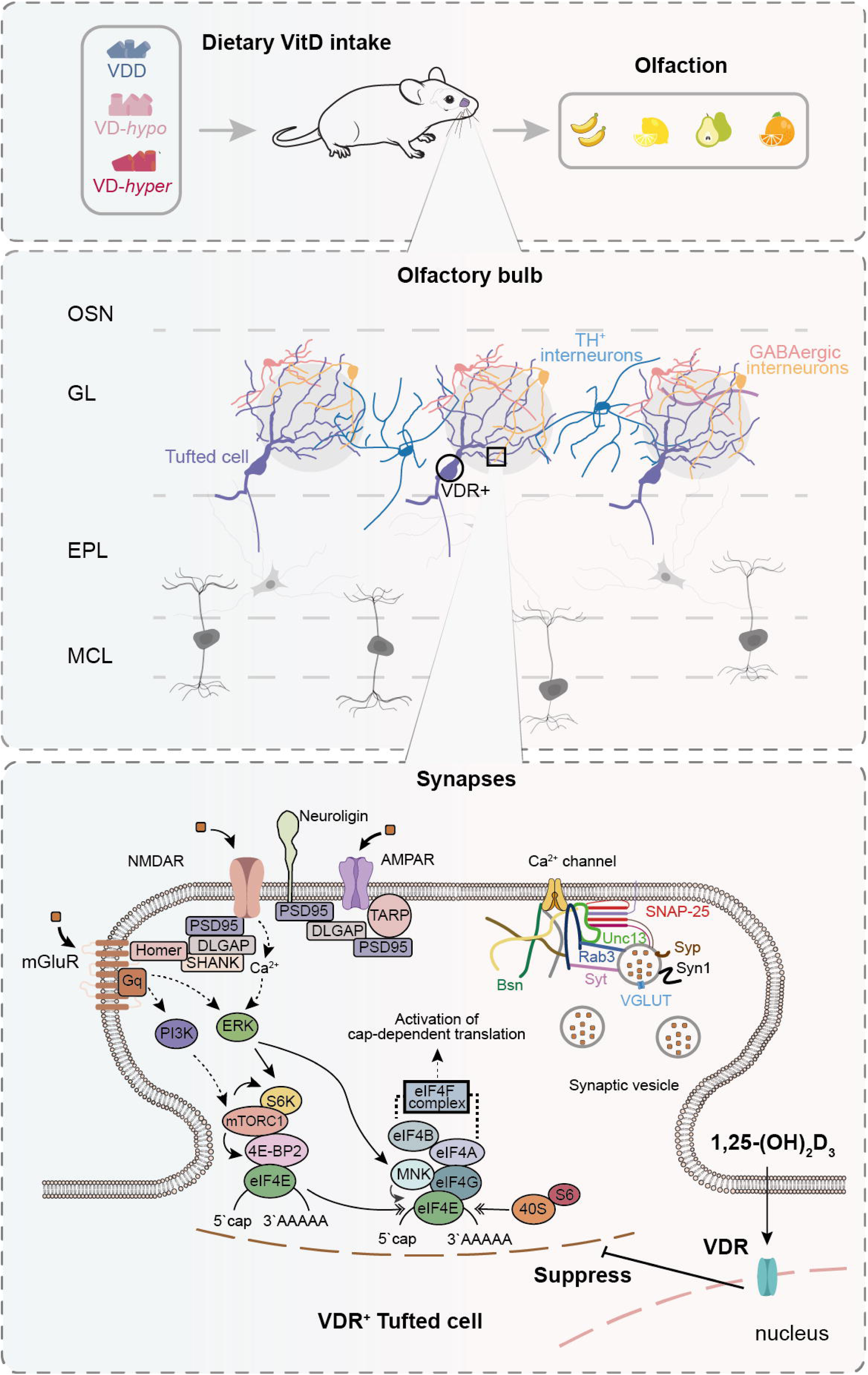

