## supplementary materials for "Vitamin D regulates olfactory function via dual transcriptional and mTOR-dependent translational control of synaptic proteins"

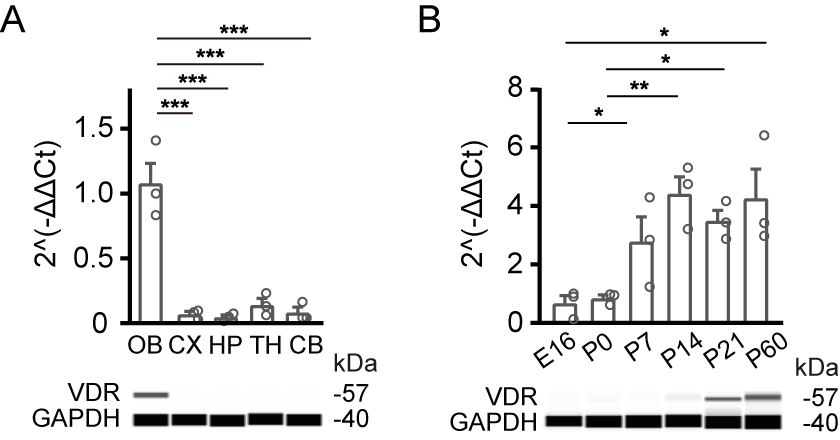

**Figure S1. VDR mRNA and protein expression in the mouse brain across various regions and developmental ages.**

**A.** Relative expression levels of VDR mRNA across different brain regions in adult mice (upper, n=3 mice), and the example *wes* image of VDR and GAPDH protein expression (lower). Brain regions include the OB, cortex (CX), hippocampus (HP), thalamus (TH), and cerebellum (CB). **B.** VDR mRNA expression levels in the mouse OB at different developmental ages, including E16, P0, P7, P14, P21, and P60 (upper, n=3 mice), and the exmaple *wes* image of VDR and GAPDH protein expression (lower, n=1 mouse). Each symbol represents a biological replicate. Data are presented as mean ± SEM. Statistical analyses were performed using one-way ANOVA followed by Tukey’s multiple comparisons. Significance levels: *P < 0.05, **P < 0.01, ***P < 0.001, ****P < 0.0001.

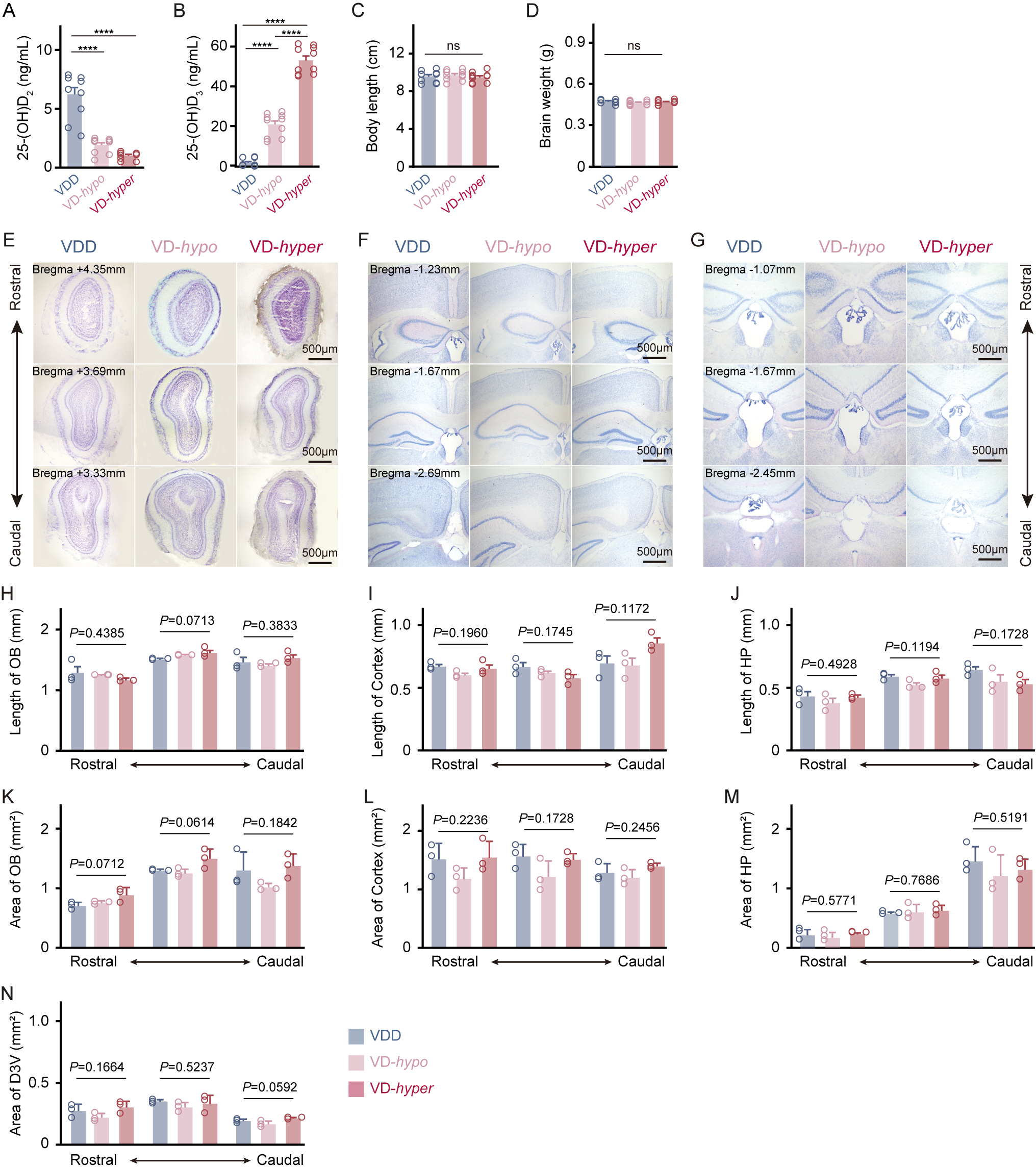

**Figure S2. Varied dietary vitamin D levels since childhood do not change brain morphology in mice.**

**A-D.** Serum levels of 25-(OH)D_2_ (**A**), 25-(OH)D_3_ (**B**), body length (**C**), and brain weight (**D**) in mice supplemented with varying doses of vitamin D3 since childhood (n=10 mice per group). **E-G.** Nissl-stained images of the OB (**E**), cortex and hippocampus (**F**), and third ventricle (**G**) from mice supplemented with different doses of vitamin D3. For each region, three coronal sections from rostral to caudal sides are presented, with corresponding bregma coordinates marked. Scale bar: 500 µm. **H-N.** Quantification of the length (**H**), and area (**K**) of the OB; length (**I**), and area (**L**) of the cortex; length (**J**), and area (**M**) of the hippocampus; and area of the third ventricle (**N**) in mice supplemented with varying doses of vitamin D3 (n=3 mice per group). Each symbol represents a biological replicate. Data are expressed as mean ± SEM and analyzed using one-way ANOVA followed by Tukey’s multiple comparisons. Significance levels: *P < 0.05, **P < 0.01, ***P < 0.001, ****P < 0.0001; ns indicates no statistical significance.

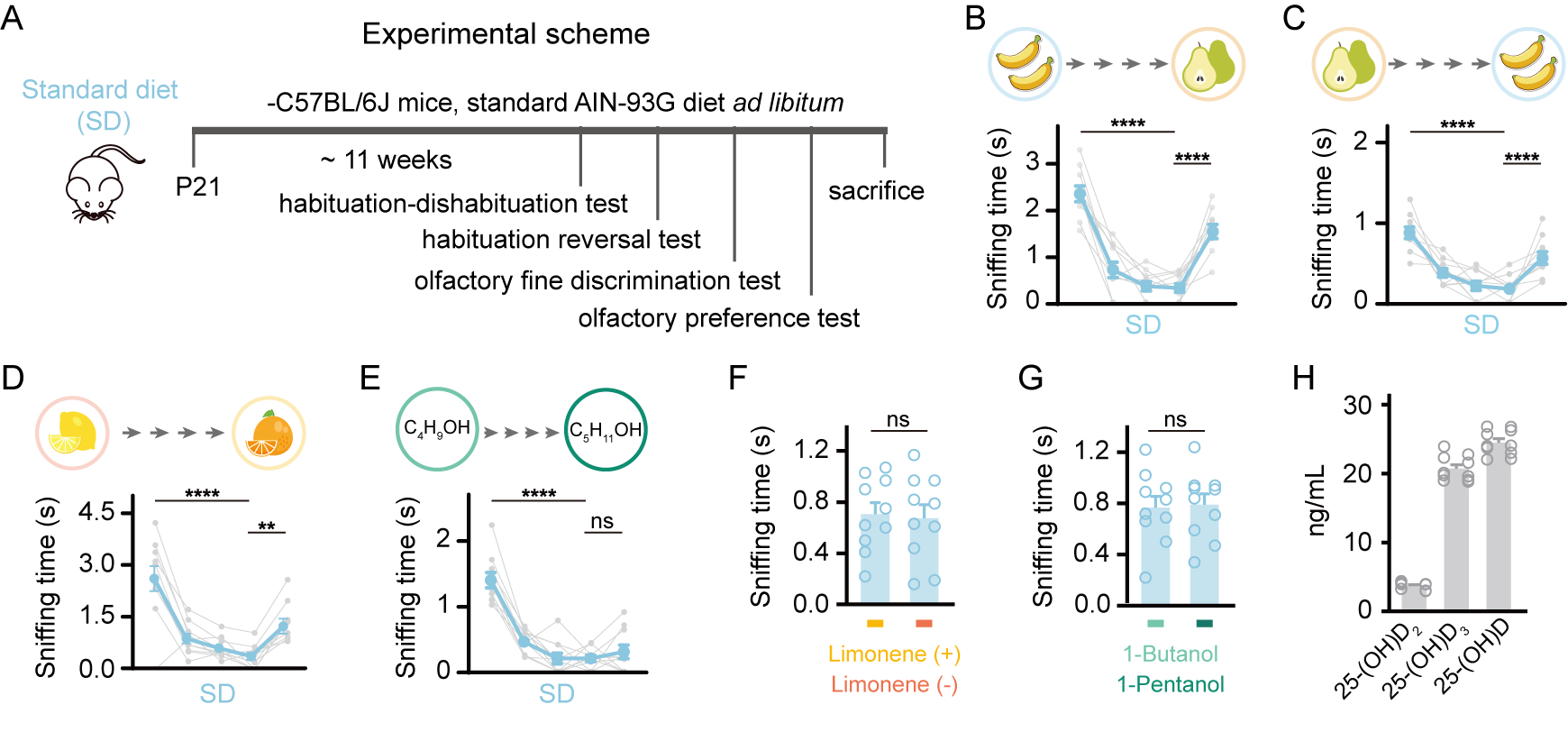

**Figure S3. Olfactory function assessment in mice fed a standard AIN-93G diet (SD group).**

**A.** Experimental timeline for behavioral testing. Mice received the AIN-93G diet *ad libitum* from P21 for 11 weeks before testing. **B-E.** Olfactory habituation-dishabituation (**B, D, E**) and habituation reversal (**C**) test results showing sniffing times (one-way ANOVA with Tukey’s correction for multiple comparisons). Tested odor pairs were: isoamyl acetate vs. 2-heptanone (**B, C**), L-limonene vs. D-limonene (**D**), and 1-butanol vs. 1-pentanol (**E**). **F-G.** Olfactory preference test results for L-limonene vs. D-limonene (**F**) and 1-butanol vs. 1-pentanol (**G**) (unpaired t-test). **H.** Serum levels of 25-(OH)D_2_, 25-(OH)D_3_, and 25-(OH)D in 18-week-old SD mice. Each symbol represents a biological replicate (n=10 mice per group). Data are presented as mean ± SEM. Significance levels: *P < 0.05, **P < 0.01, ***P < 0.001, ****P < 0.0001; ns indicates no statistical significance.

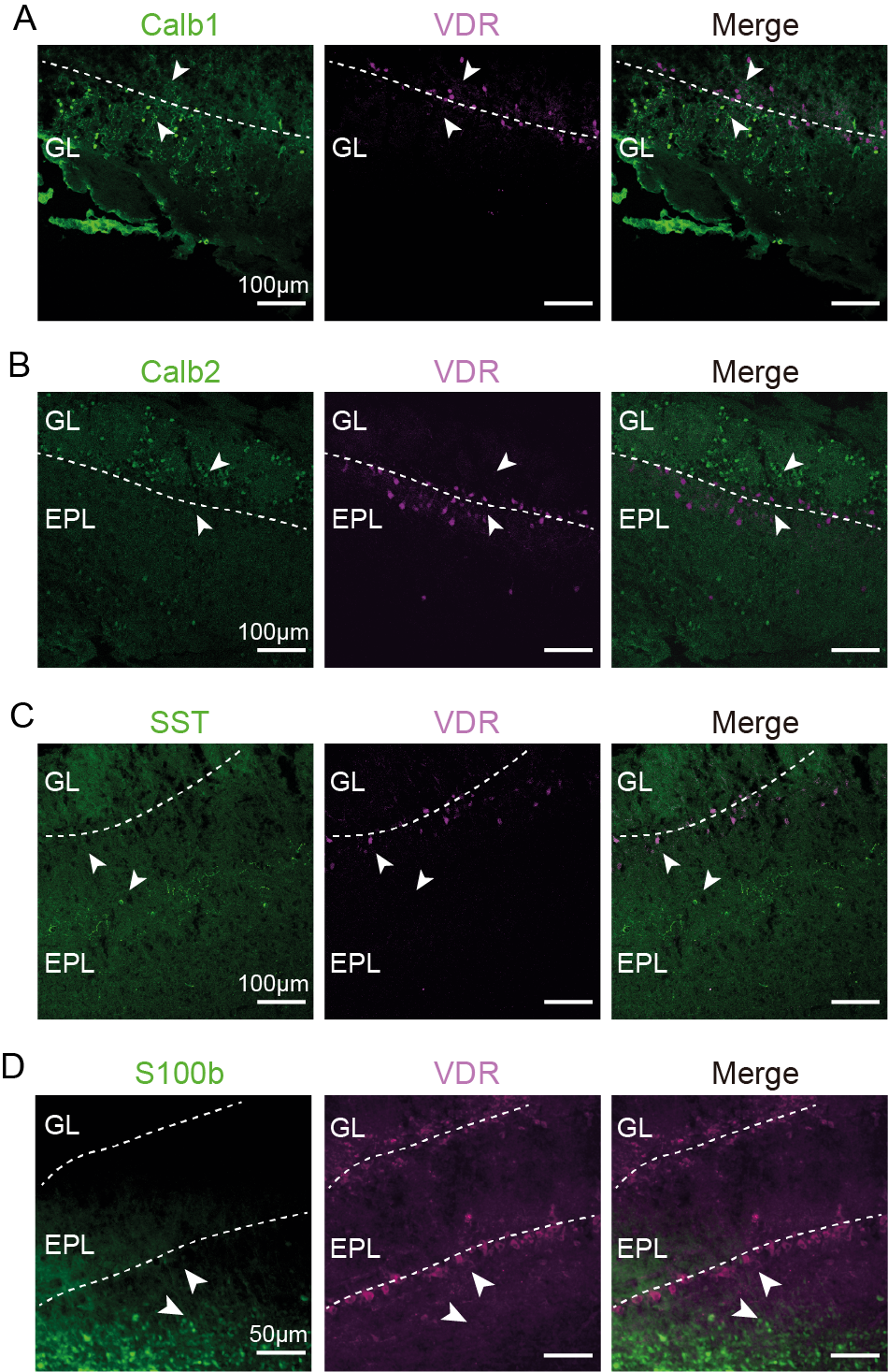

**Figure S4. Representative immunofluorescence images of VDR expression in the OB across different cell types.**

**A-D.** Representative immunohistochemistry images showing that VDR does not co-localize with Calb1 (**A**), Calb2 (**B**), SST (**C**), and S100b (**D**) in the OB (indicated by arrowheads). Purple indicates VDR staining. Scale bar: 100 µm (applies to A-C); scale bar: 50 µm (D).

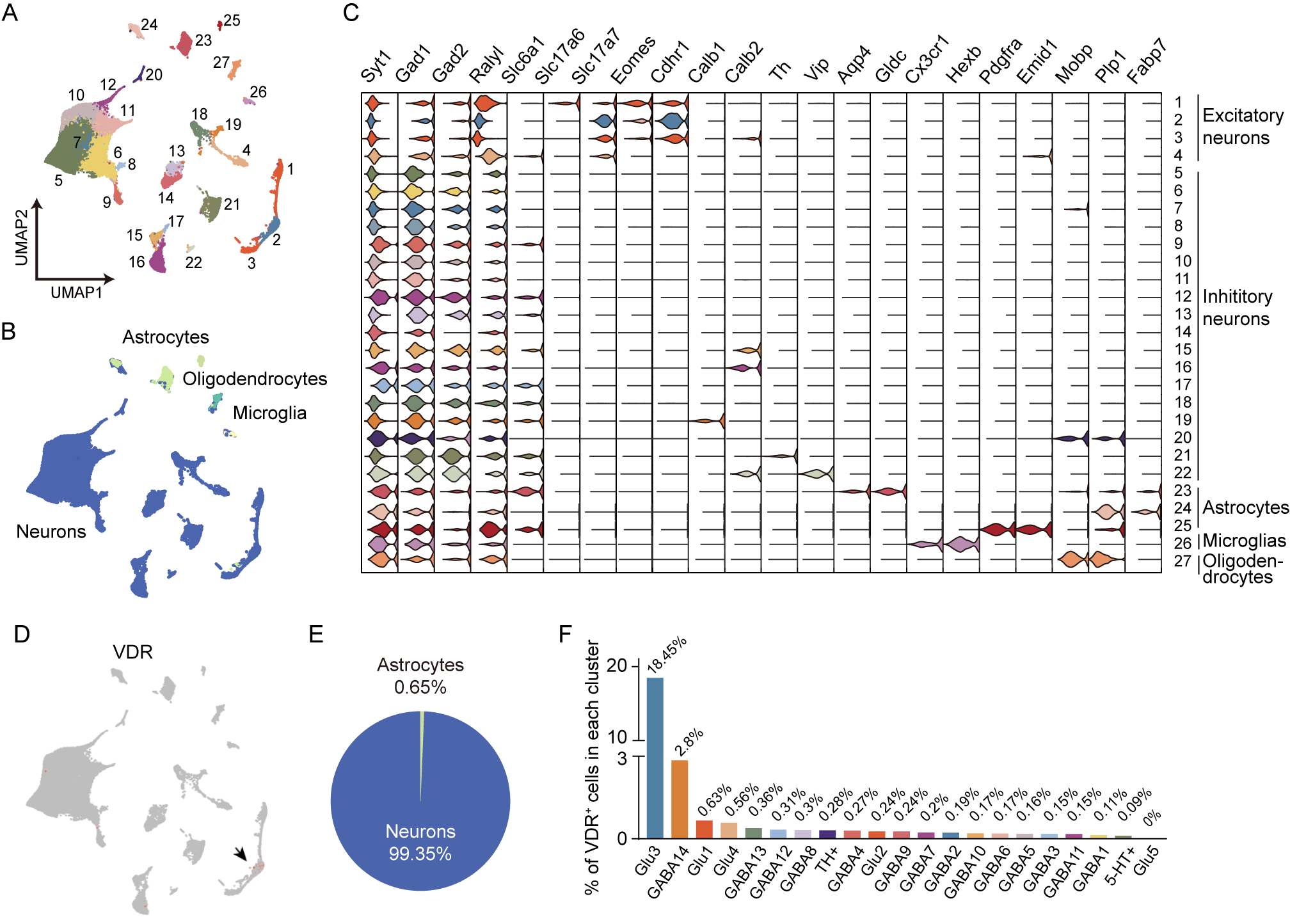

**Figure S5. snRNA-seq analysis reveals VDR expression patterns across OB cell populations.**

**A.** UMAP visualization of all cell clusters identified by snRNA-seq analysis. **B.** Cell-type classification and distribution shown by UMAP projection. **C.** Violin plots quantifying normalized expression levels of canonical cell type-specific marker genes (columns) across 27 distinct cellular clusters (rows). **D.** Spatial representation of VDR^+^ cells (red) within the neuronal and non-neuronal compartments (gray) using UMAP coordinates. The arrow head points at an excitatory neuronal cluster with high VDR expression. **E.** Quantitative cellular composition analysis demonstrating that neurons comprise 99.35% of VDR-expressing populations. **F.** Cluster-specific quantification of VDR expression frequencies (based on neuronal clusters only), with Glu3 neurons exhibiting the highest proportion (18.45%) among all populations. Data represent pooled samples from n=6 mice (2 mice from each of the three groups).

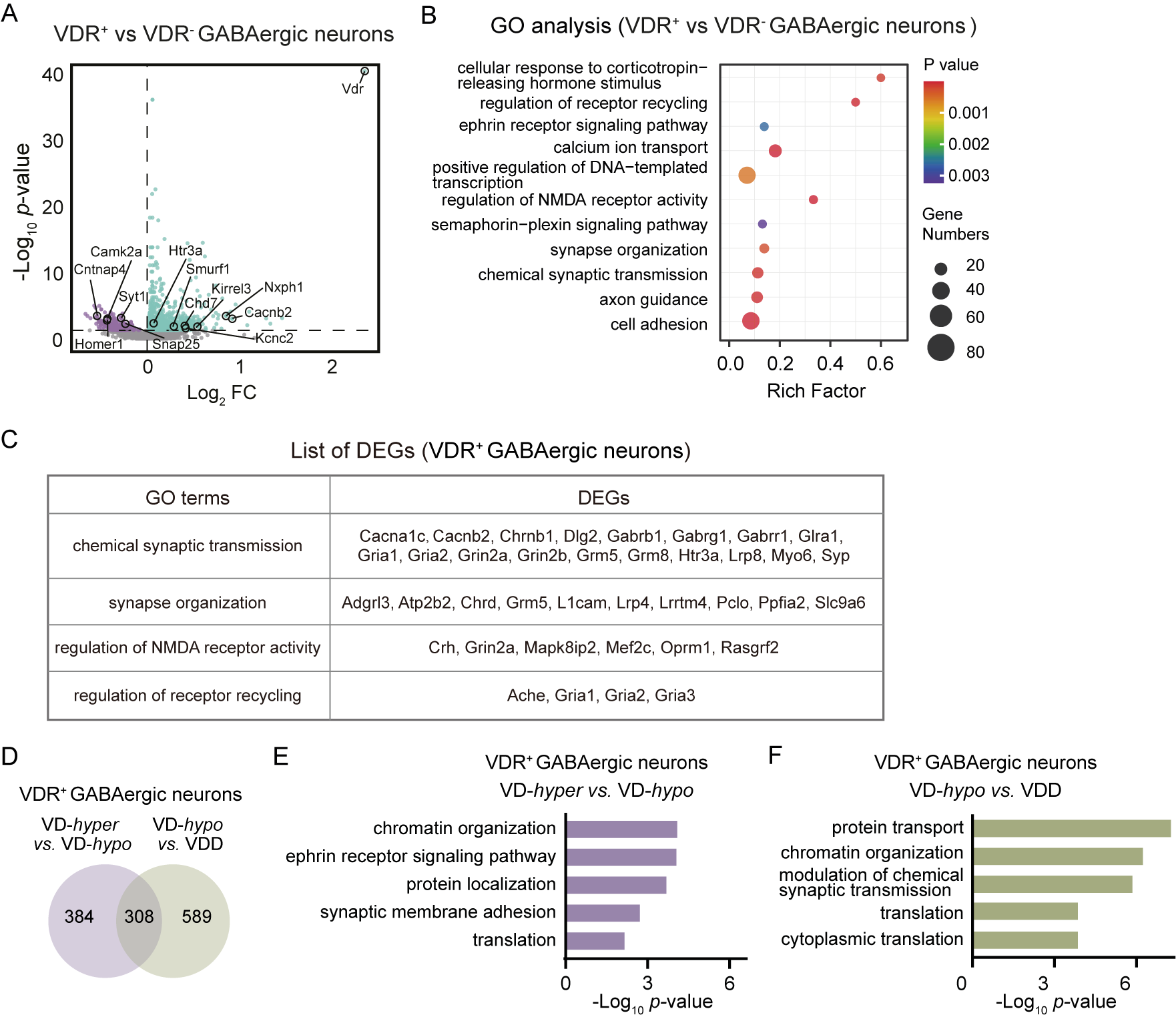

**Figure S6. DEGs and GO enrichment analysis of VDR^+^ vs. VDR^-^ GABAergic neurons in snRNA-seq.**

**A.** Volcano plot of DEGs between VDR^+^ and VDR^-^ GABAergic neurons. The x-axis represents log2-transformed fold change, and the y-axis shows statistical significance (−log10(P value)). **B.** GO annotation of DEGs between VDR^+^ and VDR^-^ GABAergic neurons. The x-axis represents the rich factor, the y-axis shows GO terms, circle size indicates the number of DEGs, and color represents the P value. **C.** List of example DEGs in selected GO terms between VDR^+^ and VDR^-^ GABAergic neurons. **D.** Venn diagram showing number of DEGs from VDR⁺ GABAergic neurons in comparisons between VD-*hyper* vs. VD-*hypo* and VD-*hypo* vs. VDD. **E-F.** GO annotation of DEGs in VDR⁺ GABAergic neurons comparing VD-*hyper* vs. VD-*hypo* (**E**) and VD-*hypo* vs. VDD (**F**). The x-axis represents −log10(P value), and the y-axis shows GO terms.

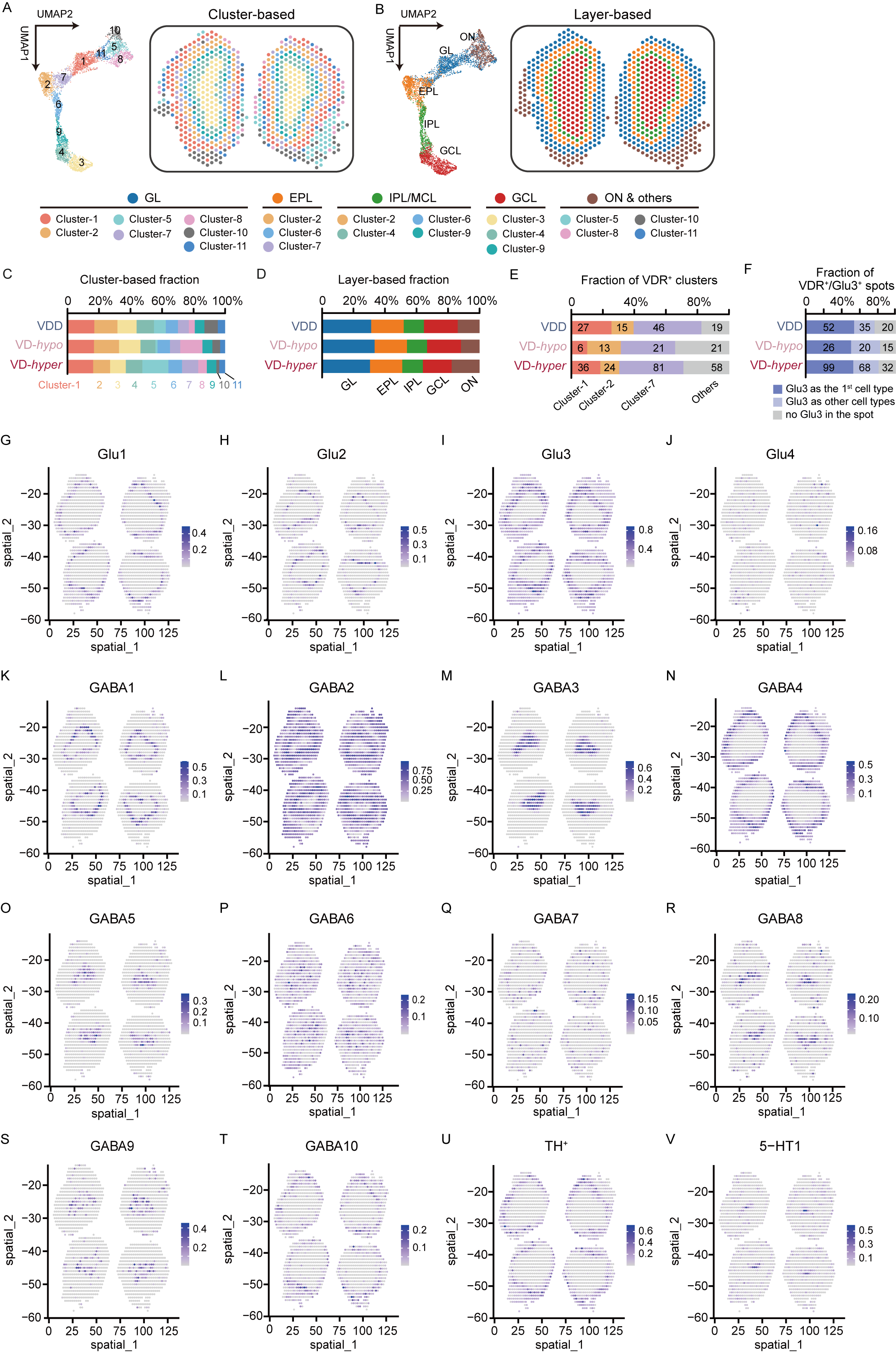

**Figure S7. Spatial transcriptomic analysis of mice fed a gradient of vitamin D3.**

**A.** UMAP plot (left) of 11 clusters classified based on spatial transcriptomic data (right; n=1 mouse per group). **B.** UMAP plot (left) of the 6 layer-based clusters classified from spatial transcriptomic data (right). **C-D.** Fraction of spots according to cluster-based (**C**) and layer-based (**D**) classification. **E.** Number and fraction of VDR^+^ spots according to cluster-based classification in three groups of mice (VDD, VD-*hypo*, and VD-*hyper*). **F.** RCTD-based classification of VDR^+^ spots into three categories: Glu3-dominant as the primary cell type, Glu3-containing as other cell type, or Glu3-absent. Stacked bars depict the category proportions of three mouse groups with total spot numbers in each case presented in the corresponding bar. **G-V.** Example images showing the distribution of individual cell types identified by RCTD analysis, including Glu1 (**G**), Glu2 (**H**), Glu3 (**I**), Glu4 (**J**), GABA1 (**K**), GABA2 (**L**), GABA3 (**M**), GABA4 (**N**), GABA5 (**O**), GABA6 (**P**), GABA7 (**Q**), GABA8 (**R**), GABA9 (**S**), GABA10 (**T**), TH^+^ (**U**), and 5-HT1 (**V**) cells.

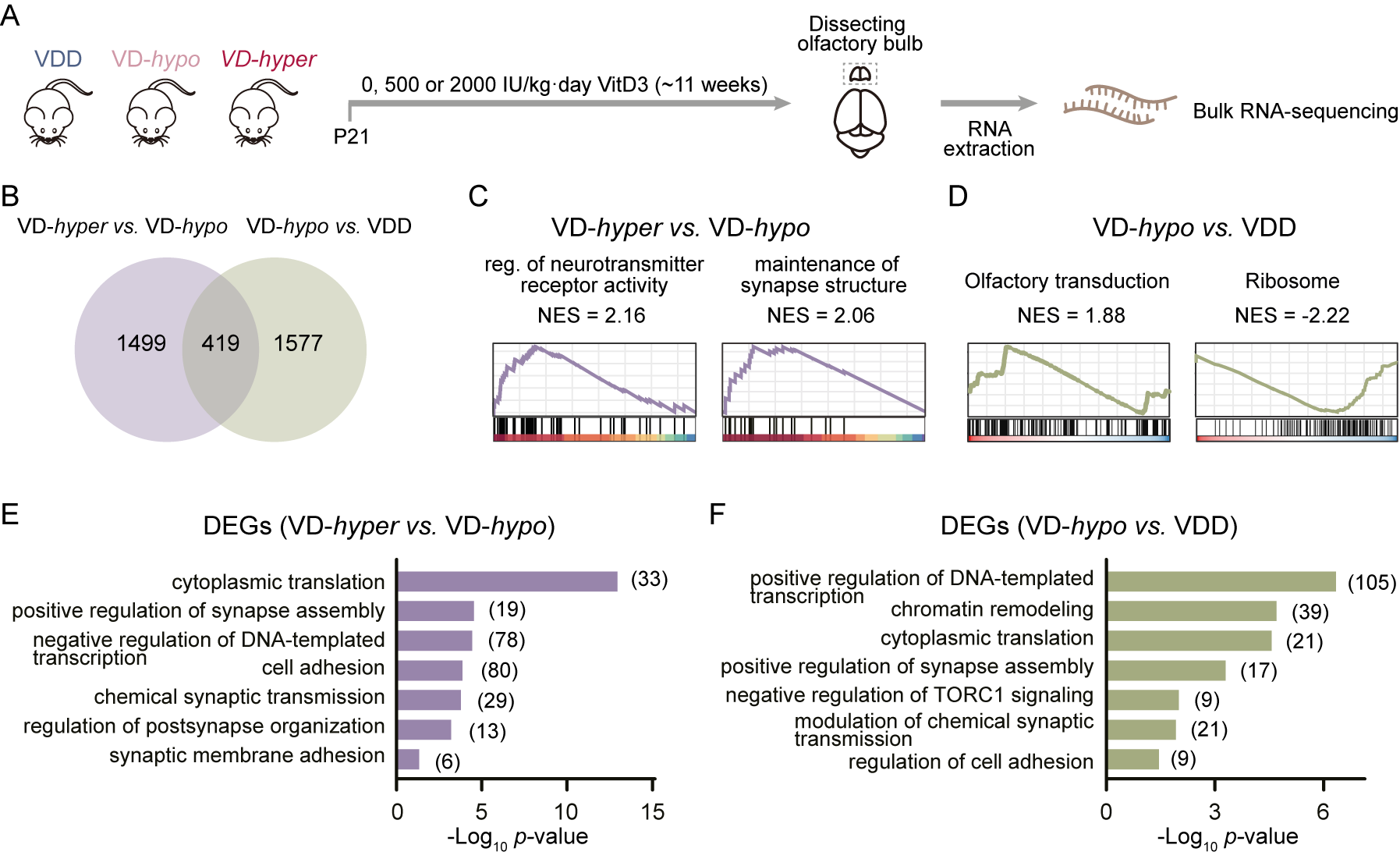

**Figure S8. Bulk RNA-sequencing reveals vitamin D3 dose-dependent pathway modulation in the OB.**

**A.** Experimental design for bulk RNA-sequencing of OBs from mice fed three vitamin D3 doses (n=3 per group). **B.** Venn diagram identifying number of DEGs between VD-*hyper* vs. VD-*hypo* and VD-*hypo* vs. VDD comparison groups. **C.** GSEA plots showing changes in neurotransmitter receptor activity (NES=2.16) and synaptic structure maintenance (NES=2.06) in VD-*hyper* vs VD-*hypo*. **D.** GSEA plots showing changes in olfactory conduction activation (NES=1.88) and ribosome suppression (NES=-2.22) in VD-*hypo* vs VDD. **E-F.** GO analysis of DEGs comparing VD-*hyper* vs. VD-*hypo* (**E**) and VD-*hypo* vs. VDD (**F**). The x-axis represents −log10(P value), and the y-axis shows GO terms. Statistical analysis conducted using standard pipelines for RNA-seq data, with normalization and multiple testing correction applied. NES values indicate normalized enrichment scores from GSEA.

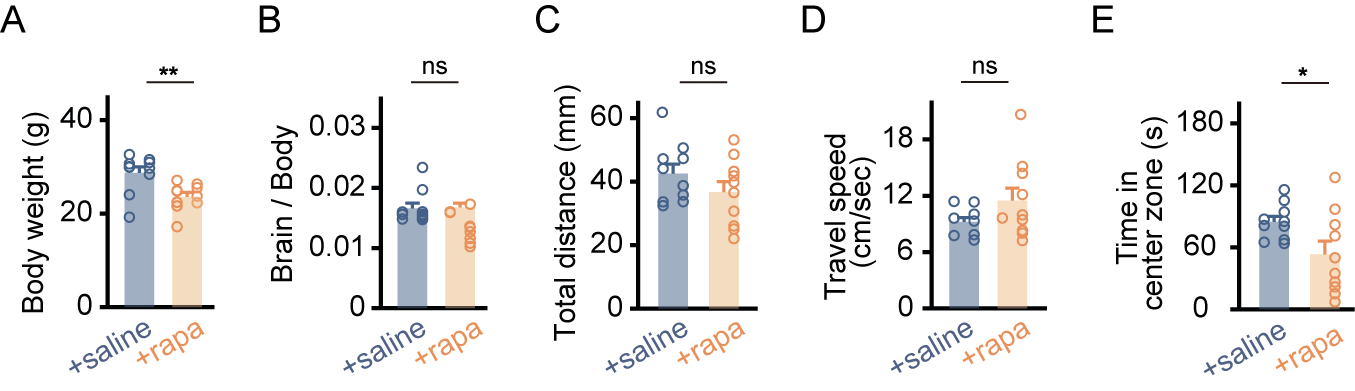

**Figure S9. Physiological and behavioral assessments of rapamycin-treated (rapa) and control (saline) mice.**

**A-B.** Body weight (**A**) and brain-to-body weight ratio (**B**) of two groups of mice (saline and rapa group). **C-E.** Total movement distance (**C**), average movement speed (**D**), time to enter the central area (**E**) of two groups of mice (saline and rapa) in the open field test. Each symbol represents a biological replicate (n=10 mice per group). All data are expressed as mean ± SEM. The comparison between two groups was analyzed using the unpaired t-test. The significance levels were defined as follows: *P < 0.05, **P < 0.01, ***P < 0.001, and ****P < 0.0001, with ns indicating no statistical significance.
